# HY5 regulates GLK transcription factors to orchestrate photomorphogenesis in *Arabidopsis thaliana*

**DOI:** 10.1101/2022.09.30.510343

**Authors:** Ting Zhang, Rui Zhang, Xi-Yu Zeng, Sanghwa Lee, Lu-Huan Ye, Shi-Long Tian, Yi-Jing Zhang, Wolfgang Busch, Wen-Bin Zhou, Xin-Guang Zhu, Peng Wang

## Abstract

Light induced de-etiolation is an important aspect of seedling photomorphogenesis. GOLDEN2 LIKE (GLK) transcriptional regulators are involved in chloroplast development, but to what extent they participate in photomorphogenesis is not clear. Here we show that ELONGATED HYPOCOTYL5 (HY5) binds to *GLK* promoters to activate their expression, and also interacts with GLK proteins. Chlorophyll content in the de-etiolating Arabidopsis seedlings of the *hy5 glk2* double mutants was lower than that in *hy5* single mutant. *GLKs* inhibited hypocotyl elongation, and the phenotype could superimpose on the *hy5* phenotype. Correspondingly, GLK2 regulates the expression of photosynthesis and cell elongation genes partially independent of HY5. Before exposed to light, the accumulation of GLK proteins was regulated by DE-ETIOLATED 1 (DET1), while also affected (especially for GLK1) transcriptionally by HY5. The enhanced etioplast development and photosystem gene expression observed in *det1* mutant were attenuated in *det1 glk2* double mutant. Our study reveals that GLKs act down-stream of HY5 and likely cooperate with HY5 and DET1, to orchestrate multiple developmental traits during the light-induced skotomorphogenesis to photomorphogenesis transition in Arabidopsis.

**One-sentence summary:** GLK and GNC act downstream of HY5, and cooperate with HY5 and DET1, to regulate both chloroplast development and hypocotyl elongation during the transition from skotomorphogenesis to photomorphogenesis.

## Introduction

Light serves as an indispensable resource and the essential regulator for plant growth. In addition to photosynthesis, light also play roles in regulating seed germination, seedling de-etiolation, organ development, flowering and other important physiological processes. After germination in darkness, seeds first grow into etiolated seedlings, displaying long hypocotyls with hooked tops and closed cotyledons. These characteristics are called skotomorphogenesis. Once the etiolated seedlings break through the soil surface and receive light, hypocotyl elongation is inhibited, cotyledons open, chloroplasts develop, and photosynthesis begins. This series of processes is referred to as photomorphogenesis. Skotomorphogenesis and photomorphogenesis together ensure that etiolated seedlings emerge from the soil and develop into normal healthy seedlings, and are both a prerequisite for plant growth and development (Sullivan and Deng, 2003; Jiao et al., 2007; Song et al., 2020).

As a positive regulator of photomorphogenesis, ELONGATED HYPOCOTYL5 (HY5) acts downstream of photoreceptors, such as cryptochromes, phytochromes and UV RESISTANCE LOCUS8 (UVR8), and plays important roles in seedling de-etiolation (Jiao et al., 2007; Stracke et al., 2010; Podolec et al., 2021; Cheng et al., 2021). Studies have shown that over 3000 genes can be directly regulated by the basic leucine zipper (bZIP) transcription factor HY5 (Jakoby et al., 2002; Lee et al., 2007; Zhang et al., 2011). It regulates many basic plant growth and development processes, including cell elongation, cell proliferation, chloroplast development, pigment accumulation, and nutrient uptake (Oyama et al., 1997; Shin et al., 2007; Jing et al., 2013; Burko et al., 2020). HY5 binds to promoters through G-box elements and inhibits the expression of cell elongation genes, such that *hy5* mutants show a typical long hypocotyl phenotype (Jing et al., 2013; Xu et al., 2016b). HY5 is important for chlorophyll and carotenoid accumulation under light as it regulates the expression of key enzymes such as PHYTOENE SYNTHASE (PSY) and PROTOCHLOROPHYLLIDE OXIDOREDUCTASE C (PORC). It also regulates the expression of photosystem proteins, such as light-harvesting chlorophyll-protein complex I subunit A4 (LHCA4) (Toledo-Ortiz et al., 2014). HY5 functions through many different regulatory interactions. For example, it interacts with HY5-HOMOLOGY (HYH) or CALMODULIN7 (CAM7) protein to bind to the HY5 promoter and activate its own expression (Kushwaha et al., 2008; Abbas et al., 2014; Binkert et al., 2014). The interaction between HY5 and B-box-containing (BBX) protein BBX21-BBX22 enhances the promoting effect of HY5 on photomorphogenesis, whereas the interaction between HY5 and BBX24-BBX25 does the opposite, suggesting that BBX protein acts as coregulator of HY5, for fine-tuning of photomorphogenesis (Datta et al., 2007; Datta et al., 2008; Xu et al., 2016a; Song et al., 2020). Furthermore, HY5 interacts with PIF1 and PIF3 proteins to coordinate the production of reactive oxygen species and the temperature control of photosynthetic gene expression (Chen et al., 2013; Toledo-ortiz et al., 2014). CONSTITUTIVE PHOTOMORPHOGENIC 1 (COP1) is a central repressor of photomorphogenesis, acting as a RING E3 ubiquitin ligase downstream of multiple photoreceptors to target key light-signaling regulators for degradation (Lau and Deng, 2012; Han et al., 2020). In the case of excess HY5 protein, it can activate the expression of COP1, thus inducing the degradation of HY5 protein and maintaining it at a relatively stable level (Huang et al., 2012).

GOLDEN2 (G2)-LIKE (GLK) and GATA NITRATE-INDUCIBLE CARBON METABOLISM-INVOLVED (GNC) are two types of transcription factors directly involved in the regulation of chloroplast development. GLK transcription factors belong to the GARP superfamily (Fitter et al., 2002), which consists of <u>G</u>LK, Arabidopsis RESPONSE REGULATOR-B (<u>AR</u>R-B) and PHOSPHATE STARVATION RESPONSE 1 (<u>P</u>SR1) from Chlamydomonas (Hall et al., 1998; Imamura et al., 1999; Wykoff et al., 1999). GLKs were first found to regulate chloroplast development in maize, with a pair of GLK homologs ZmG2 and ZmGLK1 expressed in bundle sheath and mesophyll cells respectively (Hall et al., 1998; Rossini et al., 2001). In Arabidopsis, GLK1 and GLK2 are regulated by light, and function redundantly in regulating chlorophyll biosynthesis and chloroplast development. The leaves and siliques of *glk1 glk2* double mutant are pale green, and thylakoid development in the chloroplasts is retarded (Fitter et al., 2002). Overexpression of GLK1 or GLK2 in the double mutants can restore the defect in chloroplast development (Waters et al., 2008). Ectopic expression of GLKs was also able to induce increased chloroplast numbers in non-green tissues such as roots and callus of Arabidopsis and rice (Nakamura et al., 2009; Kobayashi et al., 2012a), and in tomato fruits (Nguyen et al., 2014). GLK target genes are mainly related to chlorophyll biosynthesis, light-harvesting, and electron transport (Waters et al., 2009; Kobayashi et al., 2012b). In the process of seedling de-etiolation, GLKs interact with and are phosphorylated by BRASSINOSTEROID INSENSITIVE2 (BIN2), which promotes the stability and transcriptional activity of GLK1 (Zhang et al., 2021). GLK also interacts with ORESARA 1 (ORE1) during senescence, with ORE1 antagonizing GLK transcriptional activity (Rauf et al., 2013). The degradation of GLK itself is associated with ubiquitination process, with GLK2 shown to interact with DET1 in tomato and undergoes protein degradation mediated by the CUL4-DDB1 ubiquitin E3 ligase (Tang et al., 2016).

The *GATA NITRATE-INDUCIBLE CARBON-METABOLISM-INVOLVED* (*GNC*) gene was identified by screening mutants on nitrogen-deficient medium, with *gnc* mutants accumulating lower chlorophyll than wild type (Bi et al., 2005; Schwechheimer et al., 2022). The *CYTOKININ-RESPONSIVE GATA FACTOR1* (*CGA1*) gene, also known as *GNC-LIKE* (*GNL*), is strongly induced by cytokinin and plays a crucial role in cytokinin signaling (Naito et al., 2007). GNC and CGA1 exhibit light-dependent regulation and function redundantly in the regulation of chlorophyll biosynthesis, chloroplast division, and stomata formation (Chiang et al., 2012; Klermund et al., 2016; Ranftl et al., 2016; Bastakis et al., 2018).The *gnc cga1* double mutant exhibits decreased chlorophyll content and decreased chloroplast number. Overexpression of *GNC* or *CGA1* in Arabidopsis resulted in obvious chloroplast development in seedling roots and hypocotyls, and increased chloroplast number in cotyledon and leaf epidermal cells (Chiang et al., 2012). Overexpression of *CGA1* in rice increases chloroplast number as well as starch biosynthesis in the leaves (Hudson et al., 2013; Lee et al., 2021). GNC and CGA1 were reported to act in an additive manner with GLK1 and GLK2, to play roles in chlorophyll biosynthesis in Arabidopsis (Bastakis et al., 2018). In addition, GNC and CGA1 repress gibberellin signaling downstream from DELLA proteins and PHYTOCHROMEINTERACTING FACTORS (Richter et al., 2010). More recently, an interesting role of GNC and CGA1 in starch synthesis and gravitropic growth responses was described. It was suggested that the light-regulated GNC and CGA1 help to balance the phototropic and gravitropic growth responses during the transition from skotomorphogenesis to photomorphogenesis by inhibiting the growth of starch granules (Sala et al., 2023). Furthermore, transcriptomic studies uncovered that GNC and CGA1 are also required to repress high-light stress responses in Marchantia polymorpha and Arabidopsis thaliana (Schröder et al., 2023).

In spite of the well-known downstream targets of GLK and GNC and CGA1, their upstream regulators are relatively less established. As HY5 is a regulatory hub in light signaling, and it has been reported that GLK, GNC and CGA1 are dependent on HY5 in regulating root greening (Kobayashi et al., 2012a, 2017), the molecular relationships between HY5, GLK, GNC and CGA1 during seedling development are in need of systematic analysis. It is also important to consider the role of the COP/DET/FUS complex in regulating GLK, GNC and CGA1 stabilities. DET1 is involved in the ubiquitination and degradation of positive regulators of photomorphogenesis in darkness (Gangappa et al., 2017). Apart from shorter hypocotyls and opened cotyledons, the dark-germinated *det1* mutant seedlings show part of the light-grown traits, such as expression of photosynthetic genes and development of thylakoid membranes (Chory et al., 1989). The molecular mechanism behind such phenotypes in the dark is not fully understood and the potential involvement of GLK, GNC and CGA1is worth considering.

Therefore, this study focused on the molecular and functional linkages among HY5, GLK, GNC and CGA1 transcription factors, aiming to analyze and clarify the local regulatory network connecting them, so as to better understand the relationships between light signaling, seedling formation, and chloroplast development during photomorphogenesis. At the same time, relationships between GLK, GNC, CGA1, and DET1, and their effects on the expression of photosynthetic genes in etiolated seedlings were examined. The results shed light on the concerted involvement of HY5, GLK, GNC and CGA1 transcription factors in both skotomorphogenesis and photomorphogenesis of Arabidopsis seedlings.

## Results

### GLK2 is an effective regulator of chlorophyll content in de-etiolating Arabidopsis seedlings

HY5, GLKs, GNC and CGA1 have been reported to regulate the expression of chlorophyll biosynthesis and photosynthesis related genes (Fitter et al., 2002; Lee et al., 2007; Waters et al., 2009; Chiang et al., 2012; Hudson et al., 2013; Toledo-ortiz et al., 2014; Bastakis et al., 2018; Schröder et al., 2023). To explore the significance of these regulations during photomorphogenesis, we performed systematic investigation in de-etiolating seedlings. All plants were grown in darkness for 4 days, and then sampled at different time points in white light to determine chlorophyll content. It was found that the chlorophyll content in *glk1* mutant had less difference from the wild type than that between *glk2* mutant and the wild type (**Figures 1A and 1B; Supplemental Figure S1)**. Overexpression of GLK1 or GLK2 in *glk1 glk2* double mutants can compensate for the phenotype of decreased chlorophyll, and the chlorophyll content restored to higher level in *35S:GLK2* overexpressed material **(Supplemental Figure S1)**. These indicate that both GLKs affect chlorophyll biosynthesis in seedling de-etiolation process and GLK2 may play a dominant role. The reduced chlorophyll content in *hy5* mutant across different batches of measurements was also notable, and overexpression of HY5 in the mutant could effectively compensate for the phenotype (**Figures 1A, 1B, 9A and 9B; Supplemental Figures S1 and S2)**. We checked the expression of *GLK1, GLK2, GNC*, and *CGA1* genes in the de-etiolation process, and found that the transcript level of *GLK2* was more prominent in response to light (**Figure 1C and 1D**).

**Figure 1.**
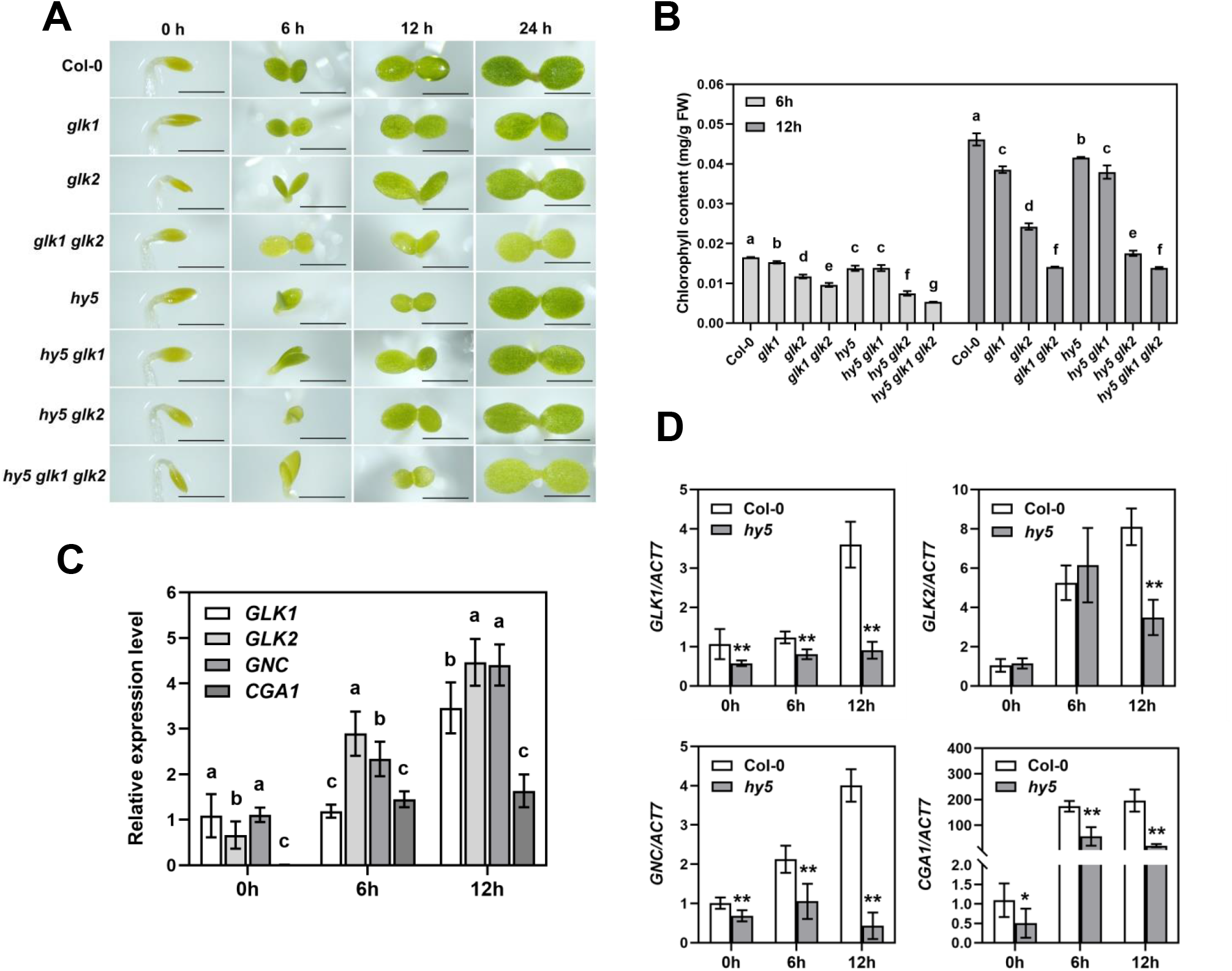
GLK2 positively regulates chlorophyll content downstream of HY5 during de-etiolation. **(A)** Phenotypes of 4-day-old etiolated seedlings of Col-0, *glk1, glk2, glk1 glk2, hy5, hy5 glk1, hy5 glk2*, and *hy5 glk1 glk2* during transition from dark to light (100 μmol/m^2^/s) conditions for 6 h, 12 h, and 24 h. Scale bars = 1 mm. **(B)** The whole seedling chlorophyll contents of de-etiolating seedlings during transition from dark to light (100 μmol/m^2^/s) for 6 h and 12 h. The data represent means ± SD (n = 3) and letters above the bars indicate significant differences (*P* < 0.05), as determined by one-way ANOVA with Turkey’s HSD test. **(C)** The relative transcript levels of *GLK1, GLK2, GNC* and *CGA1* in 4-d-old dark-grown Col-0 upon being transferred to white light at indicated time points. The value for *GLK1* at 0 h was set as “1”. The *ACT7* gene was used as internal control. The data represent means ± SD (n = 3) and letters above the bars indicate significant differences (*P* < 0.05), as determined by one-way ANOVA with Turkey’s HSD test. **(D)** The transcript levels of *GLK1, GLK2, GNC* and *CGA1* in 4-d-old dark-grown Col-0 and *hy5* seedlings upon being transferred to white light at indicated time points. The data represent means ± SD (n = 3), and asterisks indicate a significant difference compared with Col-0 (\**P*<0.05, \*\**P*<0.01, paired samples *t*-test).

### Loss of *HY5* on top of *GLK2* mutation has limited further effects on seedling greening

In order to elucidate the genetic relationships between HY5 and GLKs, we constructed double and triple mutants, and investigated the chlorophyll content in seedlings during de-etiolation. The chlorophyll content of *hy5 glk2* double mutant was lower than that of *glk2* or *hy5* mutant during greening. However, the difference in chlorophyll content of *hy5 glk1* double mutant from that of *glk1* or *hy5* mutant was less notable (**Figures 1A and 1B**).

To clarify the chlorophyll phenotype of *hy5* and *glk2* mutants, we verified the transcript levels of some chlorophyll biosynthesis and photosystem genes in de-etiolating seedlings. At 12 h of light exposure, the transcript levels of chlorophyll biosynthesis genes (*HEMA1, PORB, PORC* and *CAO*) in *hy5, glk2* and *hy5 glk2* mutants were all remarkably lower than those in the wild type. The decrease in *glk2* and *hy5 glk2* mutants was not lower than that in the *hy5* mutant, except for *PORB* (**Figure 2A**). The transcript levels of photosystem light harvesting genes (*LHCB1.2, LHCB2.2, LHCA1* and *LHCA4*) were also significantly lower in *hy5, glk2*, and *hy5 glk2* mutants than in the wild type, with those in *hy5* mutant less decreased than those in *glk2* and *hy5 glk2* mutants (**Figure 2B**). HEMA1 is a key enzyme of early chlorophyll biosynthesis in the dark, PORB and PORC are key enzymes catalyzing the light dependent transformation of protochlorophylide to chlorophylide, and CAO is responsible for switching chlorophyll a into chlorophyll b. For LHCB1.2, LHCB2.2, LHCA1 and LHCA4, they are here as representatives of the components of light-harvesting chlorophyll-binding proteins, the assembly and function of which are import for the seedling greening process. These results indicate that during de-etiolation of seedlings GLK2 has stronger regulation on photosystem light harvesting genes than HY5, which was also reflected by **Figure 5D**, whereas their regulation on chlorophyll biosynthesis may be comparable.

**Figure 2.**
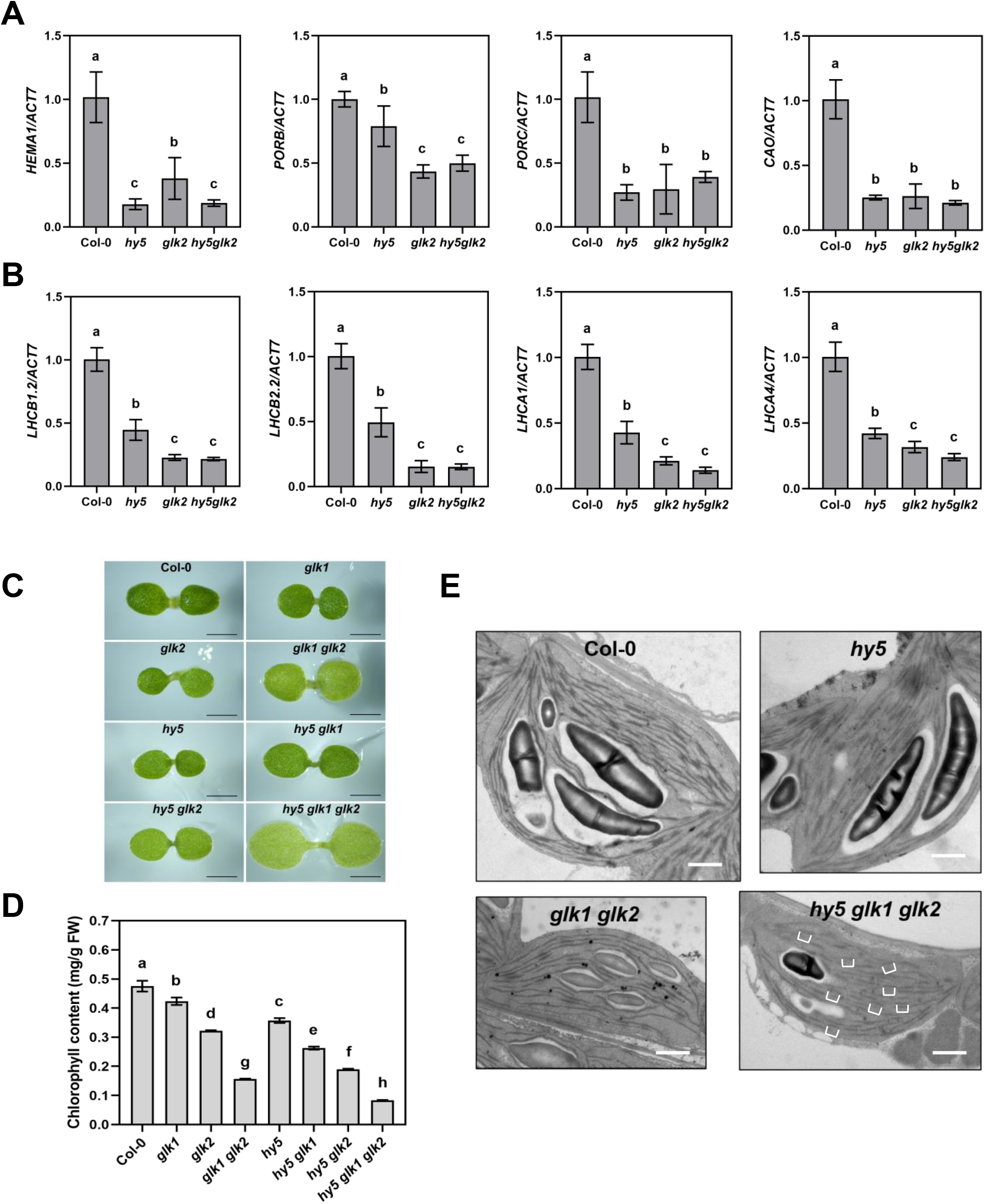
Photosynthetic gene expression and chloroplast development are further impaired in *hy5* and *glk* multiple mutants. **(A, B)** RT-qPCR analysis of chlorophyll biosynthesis and photosystem genes expressed in 4-d-old etiolated seedlings of Col-0, *hy5, glk2* and *hy5 glk2* upon being transferred to white light (100 μmol/m^2^/s) for 12 h. The *ACT7* gene was used as internal control. The data represent means ± SD (n = 3) and letters above the bars indicate significant differences (*P* < 0.05), as determined by one-way ANOVA with Turkey’s HSD test. **(C)** Phenotypes of 4-day-old Col-0, *glk1, glk2, glk1 glk2, hy5, hy5 glk1, hy5 glk2*, and *hy5 glk1 glk2* seedlings grown on half-strength MS plates under continuous white light conditions (100 μmol/m^2^/s). Scale bars = 1 mm. **(D)** Chlorophyll contents of whole seedling from the genotypes shown in (C). The data represent means ± SD (n = 3) and letters above the bars indicate significant differences (*P* < 0.05), as determined by one-way ANOVA with Turkey’s HSD test. **(E)** The ultrastructure of mesophyll cell chloroplasts of Col-0, *hy5, glk1 glk2*, and *hy5 glk1 glk2* seedlings grown under continuous white light for 7 days. The white brackets indicate some of the fractured thylakoids that are smaller in size. Scale bar = 1 μm.

To further complement the observed phenotype, we moved on to analyze the seedlings grown under continuous white light for 4 days, and found similar differences of chlorophyll content to that during de-etiolation (**Figures 2C and 2D**). We investigated the chloroplast development in wild type, *glk1 glk2, hy5*, and *hy5 glk1 glk2* mutants under continuous light exposure. Transmission electron microscopy showed that the chloroplast size of *glk1 glk2* and *hy5 glk1 glk2* mutants was smaller than that of the wild type and *hy5* mutant, the accumulation of starch grains was decreased, and the thylakoid stacking was sparser. Even worse, *hy5 glk1 glk2* triple mutant showed obvious fractured stromal thylakoids (**Figure 2E**), suggesting that *HY5* mutation combined with *GLK1* and *GLK2* double mutations can lead to further defects in chloroplast development.

### HY5 directly regulates *GLK* gene expression

As an extension of the results shown in **Figure 1C**, we found that the transcript levels of *GLK1, GLK2, GNC* and *CGA1* genes in *hy5* mutants were significantly lower than those in the wild type (**Figure 1D**), indicating that HY5 positively regulates their expression. Typical *cis*-elements to which HY5 binds to regulate target gene expression include G-box (CACGTG), T/G-box (CACGTT), E-box (CAATTG), ACE-Box (ACGT), and Z-Box (ATACGGT) (Shin et al., 2007; Chang et al., 2008; Catalá et al., 2011; Abbas et al., 2014). We performed promoter analysis for *GLK1* and *GLK2*, and found that their promoter regions contained HY5-binding *cis*-elements. These regions were synthesized to conduct electrophoretic mobility shift assays (EMSA). It was shown that HY5 could bind to *GLK1* and *GLK2* promoter sequences, competitive probes weakened such binding, while the binding ability of mutant probes was much lower than that of normal probes (**Figure 3A**). In parallel, by transforming protoplasts dual-luciferase assay displayed that HY5 could activate *GLK1* and *GLK2* expression (**Figures 3B and 3D**). While obtaining the above *in vitro* results, we also validated the binding *in vivo* via chromatin immunoprecipitation (ChIP) coupled with quantitative PCR (qPCR) assay. Three or two regions containing HY5 binding elements were selected from the promoters of *GLK1* and *GLK2* respectively, and the results demonstrated that the P2 region from *GLK1* promoter and the P1 region from *GLK2* promoter were most strongly bound by HY5 (**Figures 3C and 3E**).

**Figure 3.**
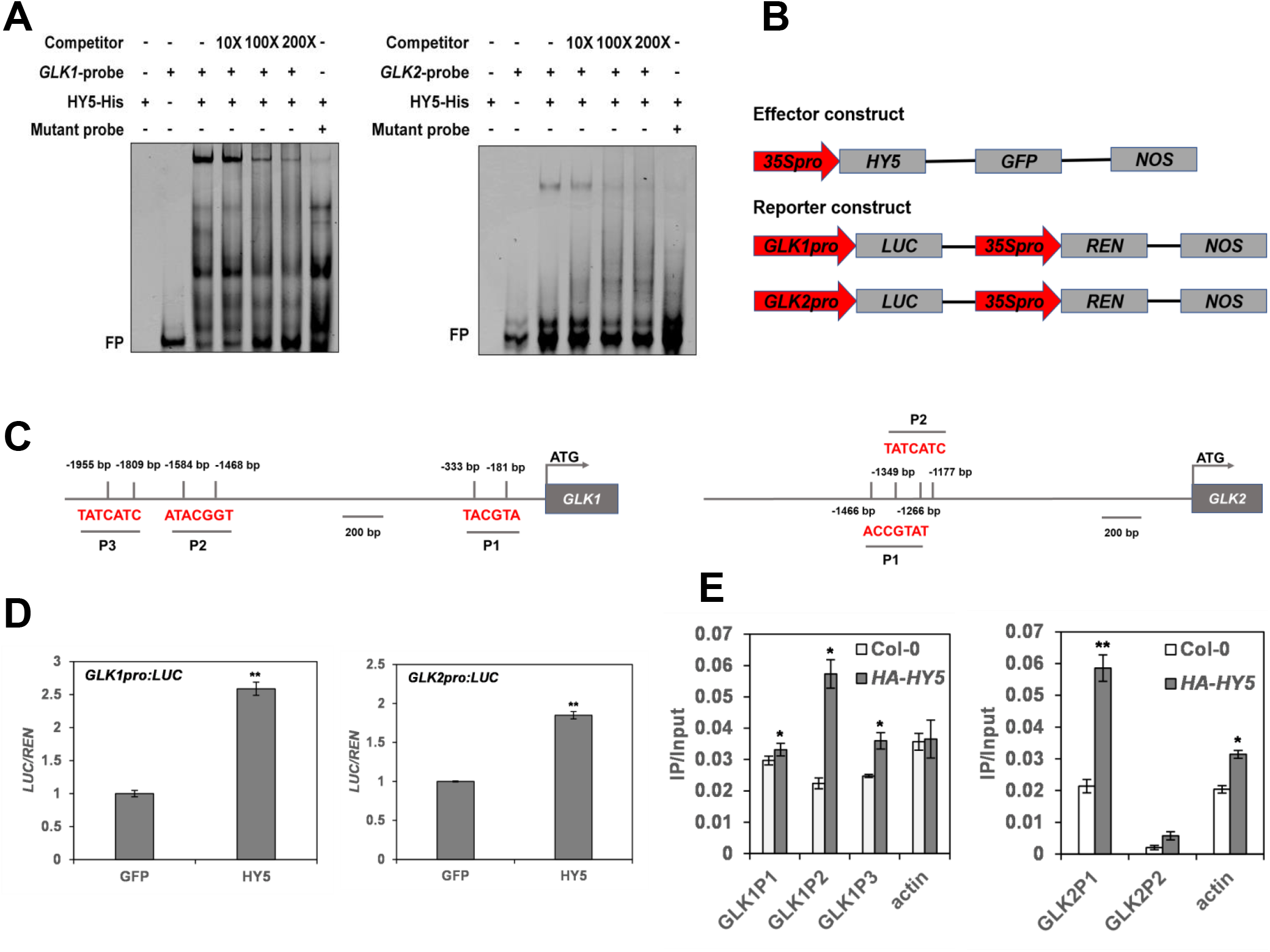
Binding of HY5 to the promoters and transcriptional activation of *GLK1* and *GLK2*. **(A)** EMSA showing HY5 binds to the sub-fragments of promoters of *GLK* and *GLK2 in vitro*. “-” and “+” indicate the absence and presence of corresponding probes or proteins. FP means free probe. **(B)** Schematic structures of effector and reporter constructs used in dual-luciferase (LUC) reporter system. *REN*, renilla luciferase gene. **(C)** Illustration of *GLK1* and *GLK2* promoter regions with the indicated positions of primers used in ChIP-qPCR. **(D)** Bar graphs showing HY5 induces the activation of *GLK1pro:LUC* and *GLK2pro:LUC*. The data represent means ± SD (n = 4), and asterisks indicate a significant difference compared with control (\**P*<0.05, \*\**P*<0.01, paired samples *t*-test). **(E)** Bar graphs showing ChIP-qPCR assays that HY5 associates with the promoters *in vivo*. Col-0 material and *ACTIN* gene were used as negative controls. The data represent means ± SD (n = 3), and asterisks indicate a significant difference compared with Col-0 (\**P*<0.05, \*\**P*<0.01, paired samples *t*-test).

### GLK negatively regulates hypocotyl elongation of light-grown seedlings

HY5 is well-known to inhibit hypocotyl elongation during photomorphogenesis, as *hy5* seedlings exhibit longer hypocotyls (Sullivan and Deng, 2003; Jiao et al., 2007). Since we found that HY5 directly regulates the expression of *GLK* genes, whether GLKs also play roles in seedling morphological development upon light regulation became the next question. Therefore, we systematically investigated the hypocotyl phenotypes of a series of *GLK* related mutants. The hypocotyl length in *glk1, glk2* and *glk1 glk2* mutants were longer than in wild type, while shorter than in *hy5* mutant, after germinating and growing under white, blue or red light for 4 days (**Figures 4A and 4B**). When *GLK1* or *GLK2* were overexpressed in *glk1 glk2* mutants, the hypocotyl length was restored to the level of wild type **(Supplemental Figure S4)**. There was no significant difference in hypocotyl length between the mutants and wild type after germinating and growing for 4 days in darkness. Together the above results indicate that GLK inhibits hypocotyl elongation during light-induced seedling development.

We further investigated the hypocotyl phenotype in *hy5* and *glk* double and triple mutants. Under white or blue light, compared with *hy5*, hypocotyls of *hy5 glk1* were slightly longer, while hypocotyls of *hy5 glk2* and *hy5 glk1 glk2* were significantly longer than that of *hy5*. The hypocotyls of all materials were generally longer under red light, and those of *hy5 glk1, hy5 glk2* and *hy5 glk1 glk2* were still longer than those of *hy5*, but with smaller differences. No notable differences in hypocotyl length were observed in dark-grown seedlings between wild type and these mutants (**Figures 4A and 4B**). We measured the cell length of wild type, *glk1 glk2, hy5*, and *hy5 glk1 glk2* materials in the middle segment of hypocotyls growing under white light for 4 days, and found that the hypocotyl cell length of *glk1 glk2* mutant was longer than that of the wild type, and that of *hy5 glk1 glk2* mutant was longer than that of *hy5* mutant (**Figures 4C and 4D**). In general, there is an obvious trend of *hy5* < *hy5 glk1* < *hy5 glk2* < *hy5 glk1 glk2* in hypocotyl length, and GLKs may work independent (at least partly) of HY5 to inhibit hypocotyl elongation, with GLK2 playing a more important role than GLK1.

The expression of several selected elongation genes was compared in wild type, *glk2, hy5*, and *hy5 glk2* mutants. Among the transcript levels of *EXPA5, IAA19, SAUR20*, and *PME16*, most of them expressed higher in *glk2* than in wild type, and higher in *hy5* than in *glk2*. They were detected higher in *hy5 glk2* than in *hy5*, with more variable degrees (**Figure 4E**). The expression of *EXPA5, SAUR20* and *PME16* were also examined in the group of wild type, *hy5, glk1 glk2*, and *hy5 glk1 glk2* mutants, and a similar ascending trend was observed **(Supplemental Figure S6)**.

**Figure 4.**
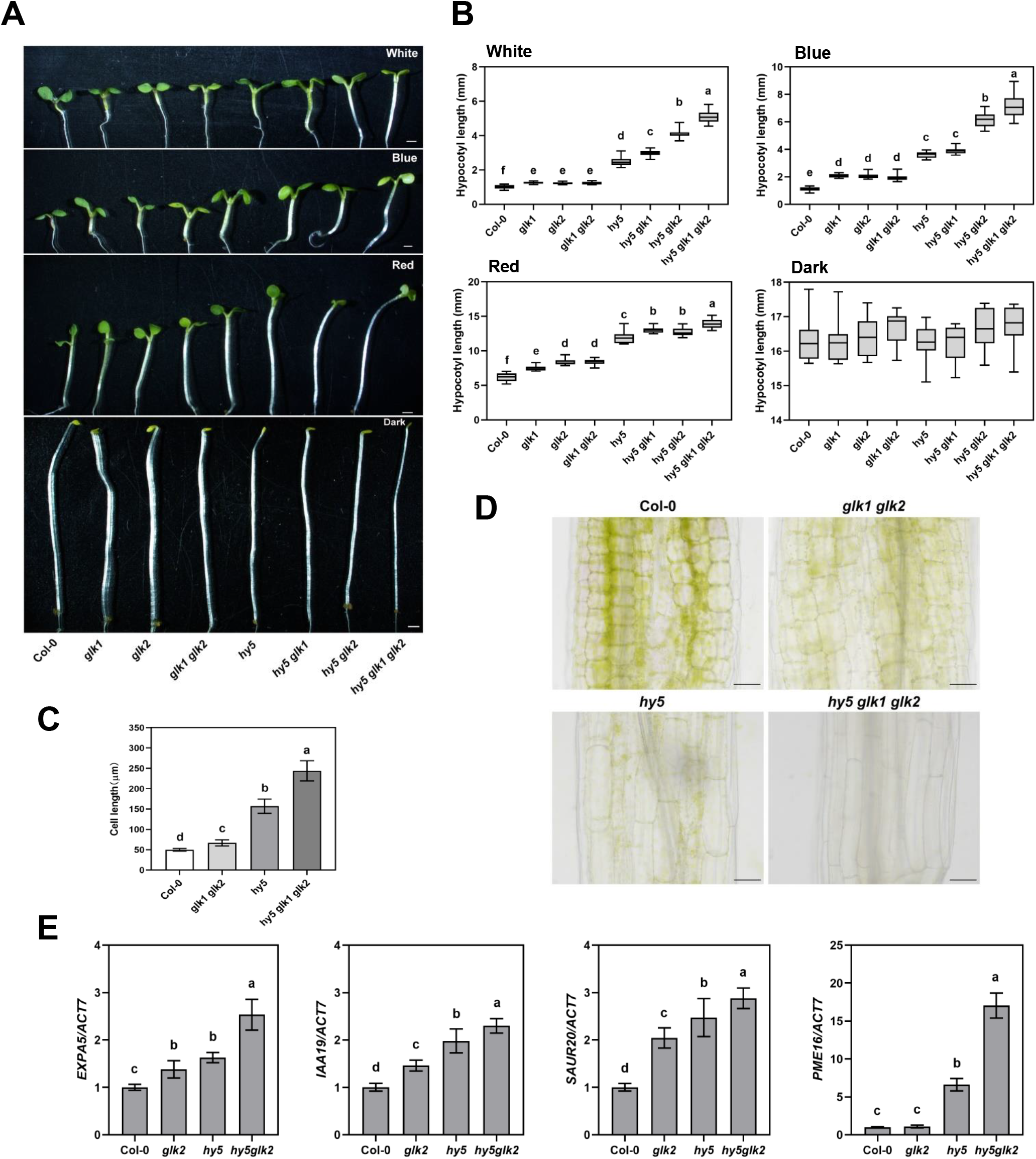
GLK1/2 negatively regulate hypocotyl elongation. **(A)** Phenotypes of 4-d-old Col-0, *glk1, glk2, glk1 glk2, hy5, hy5 glk1, hy5 glk2*, and *hy5 glk1 glk2* seedlings grown in continuous white (100 μmol/m^2^/s), blue (60 μmol/m^2^/s), red (90 μmol/m^2^/s) light and dark conditions. Scale bar = 1 mm. **(B)** Quantification of hypocotyl lengths indicated in (A). The data represent means ± SD (n ≥17) and letters above the bars indicate significant differences (*P* < 0.05), as determined by one-way ANOVA with Turkey’s HSD test. **(C)** Quantification of hypocotyl cell length from 4-d-old Col-0, *glk1 glk2, hy5*, and *hy5 glk1 glk2* seedlings grown in continuous white light (100 μmol/m^2^/s) conditions. The data represent means ± SD (n = 15) and letters above the bars indicate significant differences (*P* < 0.05), as determined by one-way ANOVA with Turkey’s HSD test. **(D)** Hypocotyl cell length phenotypes from 4-d-old Col-0, *glk1 glk2, hy5*, and *hy5 glk1 glk2* seedlings grown in continuous white light (100 μmol/m^2^/s) conditions. Scale bar = 20 μm. **(E)** RT-qPCR analysis of cell elongation genes expressed in 4-d-old Col-0, *glk2, hy5*, and *hy5 glk2* seedlings grown under continuous white light. The *ACT7* gene was used as internal control. The data represent means ± SD (n = 3) and letters above the bars indicate significant differences (*P* < 0.05), as determined by one-way ANOVA with Turkey’s HSD test.

### Transcriptome analysis reveals differential regulation of photosynthesis and cell elongation genes by HY5 and GLK2

The above results showed that the phenotype of *hy5 glk2* mutant was more severe than that of the *hy5 glk1* mutant, both in regulating chlorophyll content and inhibiting hypocotyl elongation. Hence, we selected wild type, *glk2, hy5*, and *hy5 glk2* seedlings that germinated and grew for 4 days under white light for transcriptome sequencing, in order to gain a wider spectrum of their regulated genes. As shown in a Venn diagram, *glk2, hy5*, and *hy5 glk2* mutants jointly affected the expression of 142 genes, including 138 down-regulated genes and only 2 up-regulated genes (**Figure 5A**). Deletion of *HY5* or *GLK2* in most cases resulted in down-regulation of genes, and K-means clustering showed the same trend (**Figure 5B**), in line with the expectation of them as positive regulators of photomorphogenesis. Most up-regulated genes were expressed in *hy5* and *hy5 glk2* mutants. Among the down-regulated genes, 385 were down-regulated together in *hy5* and *hy5 glk2* mutants, while 201 were down-regulated together in *glk2* and *hy5 glk2* mutants. This is consistent with HY5’s upstream position in activating *GLK* gene expression mentioned above. The remaining 91 down-regulated genes only in *hy5 glk2* mutants may represent targets of combined HY5 and GLK activity (**Figure 5A**). According to the trend of *glk2* < *hy5* < *hy5 glk2* in hypocotyl length, K4 and K7 groups from K-means clustering could be considered for further investigating GLK2 and HY5 regulated hypocotyl elongation (**Figure 5B**).

**Figure 5.**
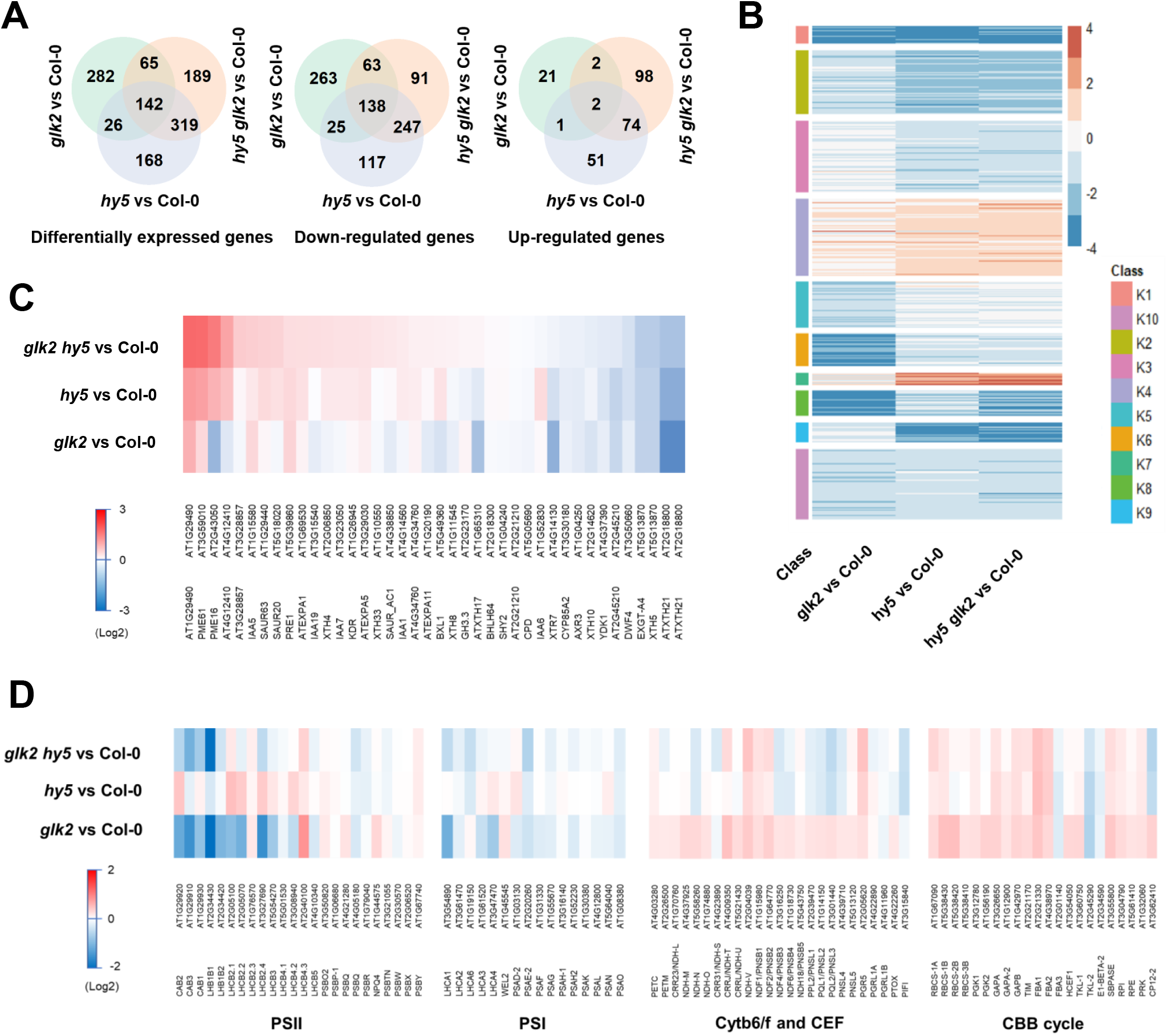
Different expression patterns of HY5 and GLK2 regulated elongation and photosynthesis genes. **(A)** Venn diagram showing overlaps between sets of differentially expressed genes in *glk2, hy5* and *hy5 glk2* mutants, relative to Col-0. **(B)** K-means clustering of differentially expressed genes regulated by GLK2, HY5 and both. The scale bar shows fold changes (log2 value). **(C, D)** Expression patterns of cell elongation genes and photosynthesis genes in *glk2, hy5* and *hy5 glk2* mutants, relative to Col-0. The scale bar shows fold changes (log2 value). Heat map was aligned according to fold changes in *hy5 glk2* mutant.

In addition, a heat map was generated from differential expression of a list of known cell elongation genes (**Figure 5C**), showing both coordinated and deviated effects on these genes by *GLK2* and *HY5* mutation. A heat map was also generated from lists of photosystem II, photosystem I, cytochrome b6f (Cyt b6f), cyclic electron transport, and Calvin-Benson-Bassham (CBB) cycle related genes, showing a differential regulation of them by HY5 and GLK2. Interestingly, GLK2 appears to positively regulate photosystem II and photosystem I, but negatively affect cyclic electron transport and CBB cycle related genes, at least at this early stage of Arabidopsis seedling development (**Figure 5D**). Whether these are direct regulation or indirect effects need further clarification.

### Interaction of HY5 and GLK proteins

Apart from directly targeting many genes to promote their expression, HY5 also interacts with other proteins, HYH or HFR1 for instance, to mutually enhance protein stability, or together bind to *HY5* or downstream gene promoters (Lee et al., 2007; Ciolfi et al., 2013; Jang et al., 2013). The interaction between HY5 and G-box binding factor 1 (GBF1) attenuates the activation of *RBCS-1A* by HY5. Since HY5 interacts with GBF1, and GBF1 interacts with GLK (Tamai et al., 2002; Singh et al., 2012; Ram et al., 2014), we then examined the possibility of HY5 interacting with GLK proteins. In pull-down experiments, the experimental group GST-GLK1 and GST-GLK2 could pull down HY5 protein, while the control group could not (**Figure 6A**), indicating that HY5 binds to GLK1 and GLK2 proteins. Through bimolecular luminescence complementation assays, it was found that luciferase signal could be detected when HY5 was co-transfected with GLK1 and GLK2 proteins, while the signal was not present in the negative controls (**Figure 6B**), indicating that HY5 interacts with GLK1 and GLK2. The above experiments provide *in vitro* evidence of HY5 and GLK interaction. We then performed co-immunoprecipitation assays to confirm that GLK1 and GLK2 interact with HY5 *in vivo*. Results showed that GLK fusion proteins were immunoprecipitated with FLAG beads, HY5 fusion protein was co-precipitated, and was detected by anti-GFP antibody (**Figures 6C and 6D**). In certain cases, HY5 might first activate the expression of *GLKs*, and then interacts with GLK proteins to regulate common targets. It should be also pointed out that extra factors might exist to coordinate the effects of HY5 and GLKs on their target genes.

**Figure 6.**
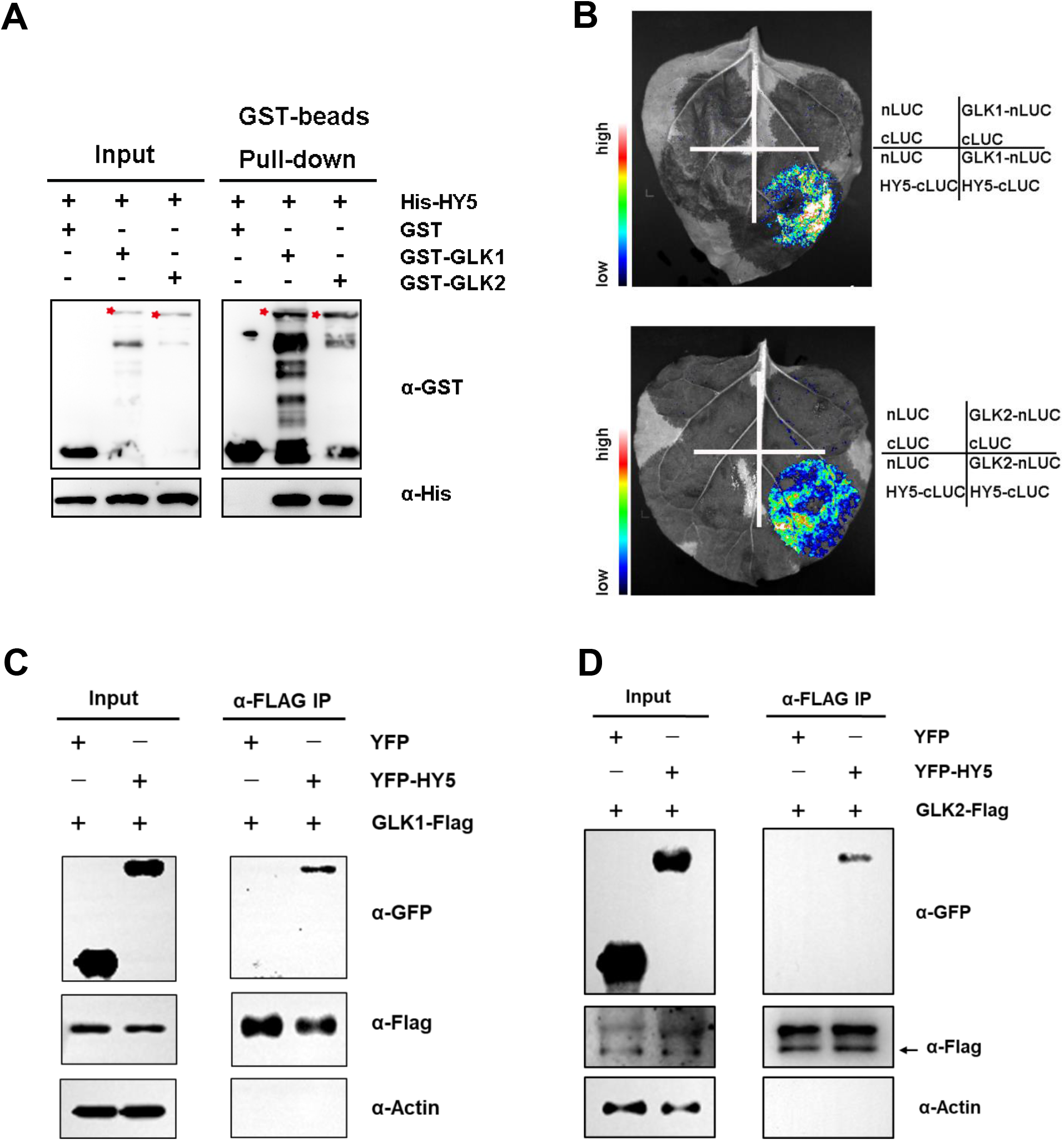
Evidences of HY5 interacting with GLK proteins. **(A)** *In vitro* pulldown assays showing the interaction of GLK1 and GLK2 with HY5. GST-GLK protein or GST protein were used to pull down His-HY5 protein using GST beads. Anti-GST and anti-His antibodies were used for immunoblot analysis. “-” and “+” indicate the absence and presence of corresponding proteins. The asterisks indicate GST-GLK1 (higher band) and GST-GLK2 (lower band) proteins. **(B)** Bimolecular luminescence complementation assays showing GLK1 and GLK2 can interact with HY5. nLUC and cLUC served as negative controls. **(C, D)** Co-immunoprecipitation assays using a transient expression system in Arabidopsis mesophyll cell protoplast, showing that GLK1 (C) and GLK2 (D) interact with HY5 *in vivo*. Protein extract was precipitated with FLAG beads, and fusion proteins were detected by immunoblot analysis using anti-GFP and anti-FLAG antibodies. The arrow indicates the band of GLK2-Flag protein.

### GLK2 content affects the plastid ultrastructure and photosystem gene expression in dark-grown *det1* mutant

In addition to the traits of photomorphogenesis, the morphology and gene expression changes of skotomorphogenesis are also points of interest for researchers. One of the well-known observations is that, the stromal thylakoid membrane was found sufficiently extended in the plastids of etiolated *det1* mutant seedlings rather than the default form of prolamellar body, and some photosynthetic genes were already expressed in the dark (Chory et al., 1989). On account of the emphasis on GLK2 from above, we constructed *det1 glk2* double mutant and comparative observations were made on the plastid ultrastructure in the cotyledons of dark-grown *det1, glk2*, and *det1 glk2* mutants. As expected, the cotyledons were open in the etiolated *det1 glk2* mutant, and paler in the light-grown *det1 glk2* mutant (**Figure 7A**). We found that the thylakoid lamellae in the plastids of *det1 glk2* mutant were less developed than in *det1* mutant in the dark, although in both cases there was little evidence of a prolamellar body, indicating that the developed chloroplasts in etiolated *det1* mutant were related to GLK2 function. By contrast, there was no significant structural difference of the prolamellar body in *glk2* mutant and *GLK2*-overexpressing line compared with the wild type, but the size was smaller or larger respectively (**Figure 7B**).

**Figure 7.**
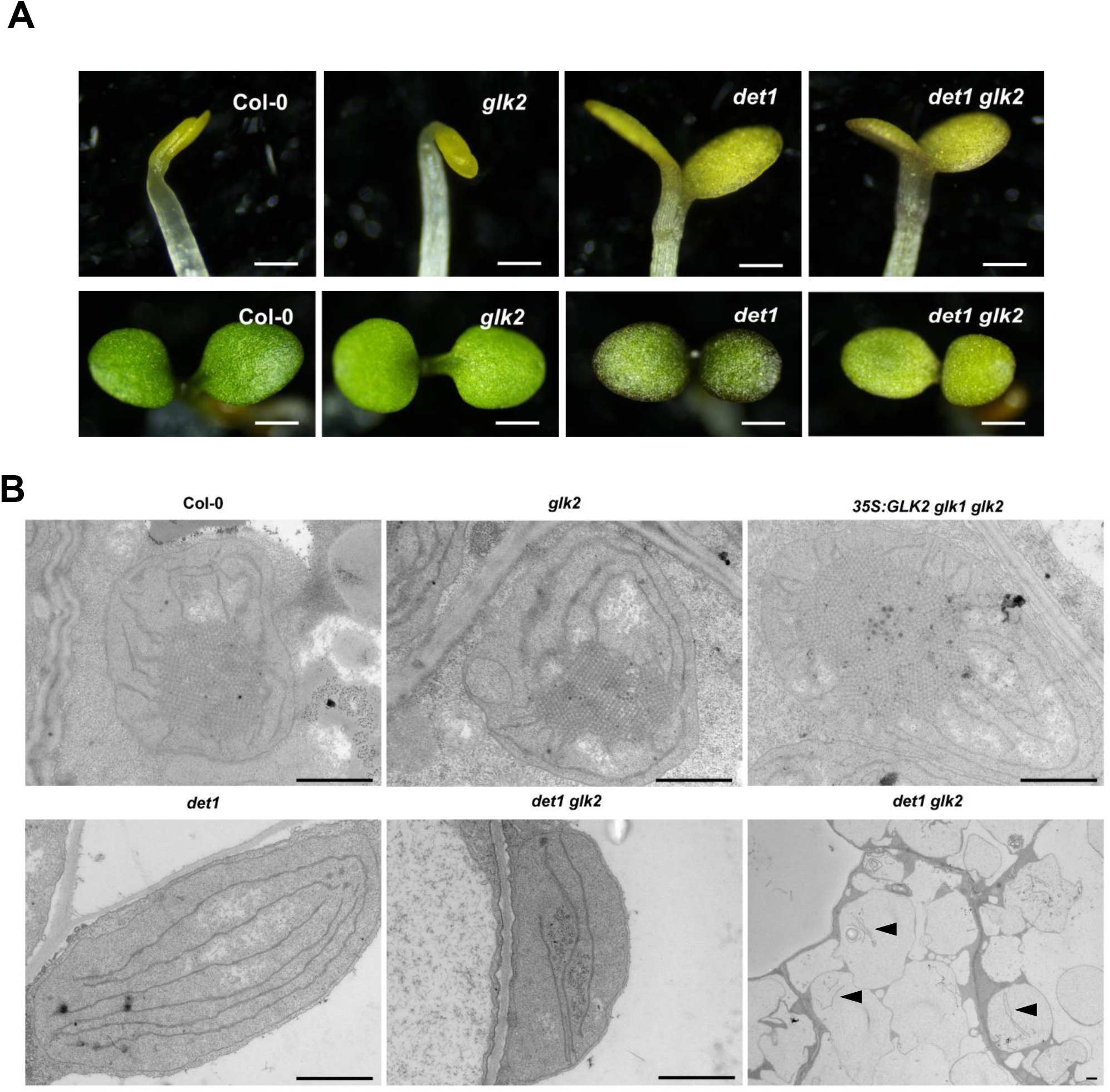
GLK2 is responsible for the developed thylakoid structures in the etiolated *det1* mutant. **(A)** Phenotype of 4-d-old Col-0, *glk2, det1*, and *det1 glk2* seedlings grown under dark (upper panel) or continuous white light (100 μmol/m^2^/s) conditions (lower panel). Scale bars represent 500 μm (upper panel) or 1 mm (lower panel). **(B)** The ultrastructure of mesophyll chloroplasts of Col-0, *glk2, 35S:GLK2 glk1 glk2, det1*, and *det1 glk2* seedlings grown under dark condition for 4 days. Scale bar = 1 μm. Black arrow heads in the bottom right image point to the thylakoid lamellae in very few amount.

It is known that in the dark, HY5 ubiquitination and degradation are mediated by the COP/DET/FUS complex (Saijo et al., 2003). In order to study the stability of GLK proteins in Arabidopsis, we examined the dark-grown wild type and *det1* mutant using Arabidopsis GLK1 and GLK2 antibodies for Western-blots. The results showed that both GLK1 and GLK2 proteins were evidently accumulated in *det1* mutant whereas they are hardly detectable in the wild type, *glk1 glk2*, and *hy5* (**Figure 8A**), suggesting that DET1 may promote the degradation of GLK proteins. It should be noted that Rubisco large subunit (bands slightly larger than 50 kD in Coomassie stained PAGE gel) content was higher in *det1* than in the wild type, which is consistent with the increased accumulation of *RbcL* transcript in dark-grown *det1* mutant (Chory et al., 1989). After MG132 (a proteasome inhibitor) treatment, the accumulation of GLK proteins in the wild type was increased, but still much lower than that in the *det1* mutant (**Figure 8A**). The deficiency of DET1 likely blocked the upstream steps of protein degradation, so that the accumulation of GLK proteins in *det1* mutant appeared more effective. Supportively, *in vitro* protein binding experiments indicate that DET1 interacts with both GLK1 and GLK2 proteins, and GNC and CGA1 proteins as well **(Supplemental Figure S9)**. On the other hand, the transcript levels of *GLK1* and *GLK2* genes were significantly higher in etiolated *det1* mutant than in etiolated wild type (**Figure 8B**), which should be considered together with the HY5 regulation of GLKs and HY5 protein stability in *det1* mutant (Saijo et al., 2003; Cañibano et al., 2021).

**Figure 8.**
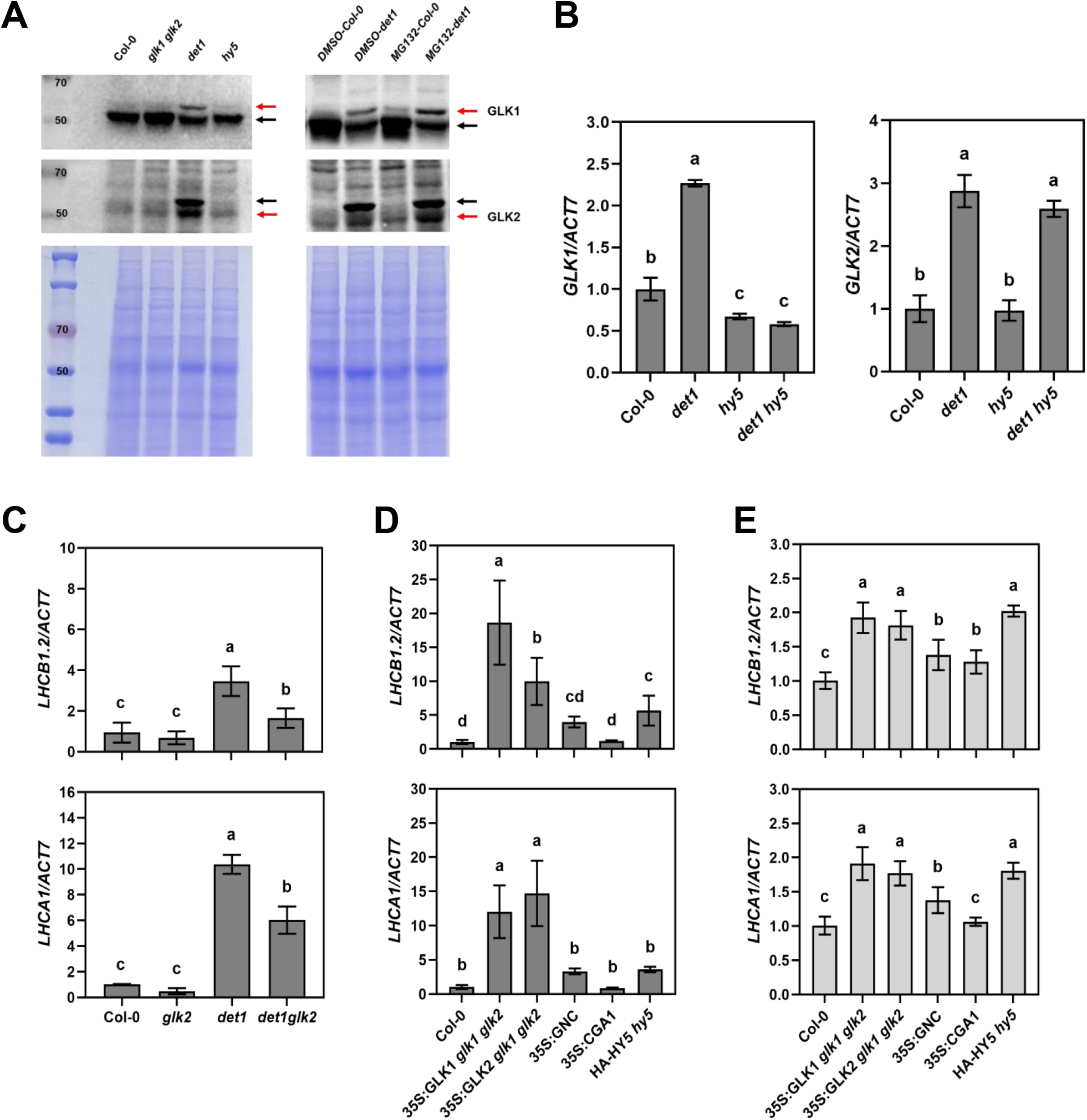
DET1 promotes the degradation of GLK proteins, and GLK2 is responsible for photosystem gene expression in the etiolated *det1* mutant. **(A)** Immunoblot detection of GLK1 and GLK2 protein level in Col-0, *glk1 glk2, det1* and *hy5* seedlings grown in the dark for 4 d (left). GLK protein levels in 4-d-old dark-grown Col-0 and *det1* seedlings after 12 h treatment with 100 μM MG132 or DMSO (right). Red arrows indicate target bands; black arrows indicate non-specific bands. Coomassie stained PAGE gel was shown as loading control. See Supplemental Figure S10 for validation of GLK2 antibody. (**B**) RT-qPCR analysis of *GLK* genes expressed in 4-d-old Col-0, *det1, hy5*, and *det1 hy5* seedlings grown under dark condition. (**C**) RT-qPCR analysis of photosystem genes expressed in 4-d-old Col-0, *glk2, det1*, and *det1 glk2* seedlings grown under dark condition. (**D, E**) RT-qPCR analysis of photosystem genes expressed in 4-d-old Col-0, *35S:GLK1 glk1 glk2, 35S:GLK2 glk1 glk2, 35S:GNC, 35S:CGA1*, and *HA-HY5 hy5* seedlings grown under dark (D) or continuous white light-grown (100 μmol/m^2^/s) (E) conditions. The *ACT7* gene was used as internal control. The data represent means ± SD (n = 3) and letters above the bars indicate significant differences (*P* < 0.05), as determined by one-way ANOVA with Turkey’s HSD test.

The expression of photosynthetic genes in dark-grown *det1* mutant seedlings is interesting (Chory et al., 1989), but the mechanism of these genes being turned off in skotomorphogenesis, has not been elucidated. We measured the expression of photosystem genes in a group of dark-grown *det1* and *glk2* related mutants, and found that the transcript levels of *LHCB1.2* and *LHCA1* were much higher in *det1* mutant, but were lower in *glk2* mutant than those in wild type. The down-regulation of these genes by GLK2 mutation was also found in *det1 glk2* double mutant comparing to *det1* mutant (**Figure 8C**). This indicates that GLK2 contributes significantly to the expression of photosystem genes in dark-grown *det1* seedlings. In the etiolated materials overexpressing *GLK1, GLK2, GNC* or *CGA1*, the expression of *LHCB1.2* and *LHCA1* genes were significantly promoted in *GLK1* or *GLK2* overexpression lines, less promoted in the *GNC* overexpression line, and least changed in the *CGA1* overexpression line (**Figure 8D**). When this same set of materials were grown in the light, increased expression of *LHCB1.2* and *LHCA1* genes were still observed to larger extent in *GLK1* or *GLK2* overexpression lines than in *GNC* and *CGA1* overexpression lines relative to the wild type (**Figure 8E**). Notably, transcript levels of *LHCB1.2* and *LHCA1* genes also increased in both dark and light-grown *HY5* overexpression lines, but were not higher than those in *GLK1* or *GLK2* overexpression lines, indicating again that GLKs function at least partly independent of HY5 during seedling photomorphogenesis.

### GNC and CGA1 positively regulate both seedling greening and hypocotyl elongation

Having obtained various clues of GLK involvement in seedling photomorphogenesis, we also scanned the set of *GNC* and *CGA1* related materials for their performance. The differences of chlorophyll contents of *GNC, CGA1* related mutants or overexpression lines compared to the wild type, appeared smaller than that between *glk2* mutant and the wild type. The chlorophyll content in *hy5 gnc* and *hy5 cga1* mutants also showed less difference compared to *hy5* during de-etiolation (**Figures 9A and 9B**). On another hand, *in vitro* experiment EMSA and dual-luciferase assays confirmed that HY5 could also bind to *GNC* and *CGA1* promoters to activate their transcription, again followed by *in vivo* experiment ChIP-qPCR showing that the highest enrichment of P1 on *GNC* and *CGA1* promoters by HY5 **(Supplemental Figure S3)**.

**Figure 9.**
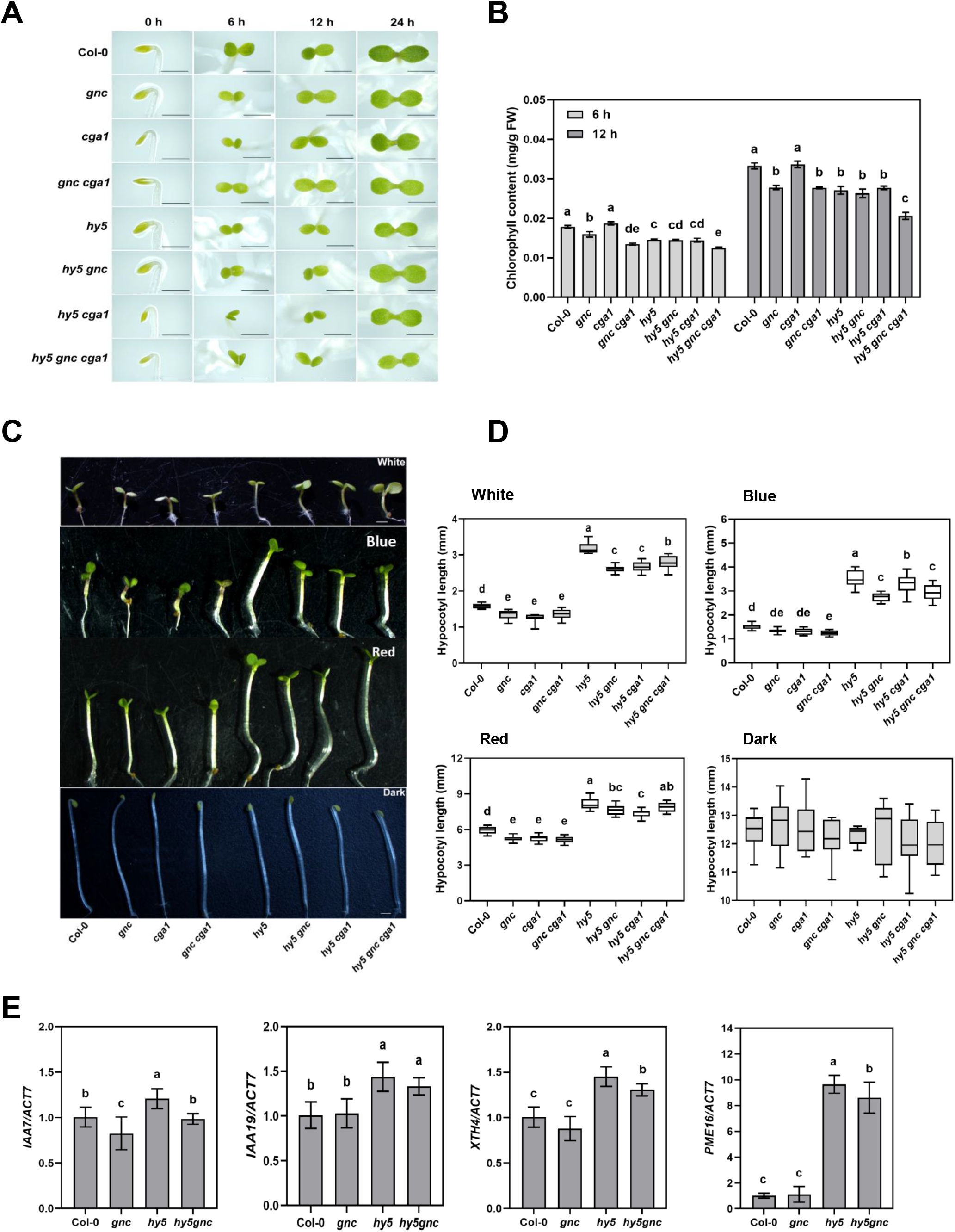
GNC and CGA1 positively regulate hypocotyl elongation. **(A)** Representative images of 4-day-old etiolated seedlings of Col-0, *gnc, cga1, gnc cga1, hy5, hy5 gnc, hy5 cga1,* and *hy5 gnc cga1* during the transition from dark to light (100 μmol/m^2^/s) conditions for 6 h, 12 h, and 24 h. Scale bars = 1 mm. **(B)** The whole seedling chlorophyll contents of 4-day-old etiolated seedlings from (A) during the transition from dark to light conditions for 6 h and 12 h. The data represent means ± SD (n=3) and letters above the bars indicate significant differences (*P* < 0.05), as determined by one-way ANOVA with Turkey’s HSD test. **(C)** Phenotypes of 4-d-old Col-0, *gnc, cga1, gnc cga1, hy5, hy5 gnc, hy5 cga1*, and *hy5 gnc cga1* seedlings grown in continuous white light (100 μmol/m^2^/s), blue (60 μmol/m^2^/s), red (90 μmol/m^2^/s) light and dark conditions. Scale bar = 1 mm. **(D)** Quantification of hypocotyl lengths indicated in (C). The data represent means ± SD (n ≥15) and letters above the bars indicate significant differences (*P* < 0.05), as determined by one-way ANOVA with Turkey’s HSD test. **(E)** RT-qPCR analysis of cell elongation genes expressed in 4-d-old Col-0, *gnc, hy5*, and *hy5 gnc* seedlings grown under continuous white light (100 μmol/m^2^/s). The *ACT7* gene was used as internal control. The data represent means ± SD (n = 3) and letters above the bars indicate significant differences (*P* < 0.05), as determined by one-way ANOVA with Turkey’s HSD test.

Different from the hypocotyl phenotype of *glk* series mutants, after germinating and growing under white, blue, or red light for 4 days, the hypocotyl length of *gnc, cga1* and *gnc cga1* mutants was shorter than that of the wild type, while the hypocotyl length of *35S:GNC* and *35S:CGA1* overexpressed materials was longer than that of the wild type. There was no difference in hypocotyl length between these mutants and wild type after germinating and growing in darkness (**Figures 9C and 9d; Supplemental Figure S5**), indicating that GNC and CGA1 promoted hypocotyl elongation under light. In addition, the hypocotyl length of *hy5 gnc, hy5 cga1* and *hy5 gnc cga1* mutants was shorter than that of *hy5*, but significantly longer than that of *gnc, cga1* and *gnc cga1* mutants (**Figures 9C and 9D**), indicating that GNC and CGA1 were able to reduce the inhibition of HY5 on hypocotyl elongation.

We also analysed the transcript levels of selected elongation genes in *gnc* and *hy5 gnc* materials. The transcript level of *IAA7* in *gnc* mutant was lower than that in wild type, and the transcript levels of several other genes in *gnc* mutant tended to be lower than in wild type, although not significant. In addition, the transcript levels of *IAA7, IAA19, XTH4*, and *PME16* appeared lower in the *hy5 gnc* double mutant than in *hy5* mutant (**Figure 9E**). These results indicate that GNC could promote the expression of these elongation genes to less and varying degrees, either independently or on top of HY5. The expression of *EXPA5, SAUR20* and *PME16* were also examined in the group of wild type, *hy5, gnc cga1*, and *hy5 gnc cga1* mutants, and a similar pattern to that with *gnc* and *hy5 gnc* was observed **(Supplemental Figure S6)**. It should be noted however, that the primarily targeted hypocotyl elongation genes by GNC and CGA1 may be largely different from those by GLKs.

## Discussion

In this work, we have constructed a series of mutant materials to systematically study the regulatory relationships between HY5, GLK, GNC and CGA1 transcription factors, as well as their stabilities and functions within the scenario of skotomorphogenesis and photomorphogenesis. We find that: (1) HY5 directly activates *GLK, GNC*, and *CGA1* expression, and together they positively regulate chlorophyll biosynthesis and photosystem formation; (2) GLK proteins inhibit hypocotyl elongation, whereas GNC and CGA1 promote hypocotyl elongation, by differentially regulating the expression of cell elongation genes; (3) In the dark, the accumulation of GLK proteins was likely both HY5 and DET1 dependent, and the abnormal light-grown traits exhibited in the etioplast of *det1* mutant seedlings can be attributed to GLK2 activity; (4) GLK2 maybe the predominant regulator of photosynthetic genes, during both skotomorphogenesis and photomorphogenesis. At present, the molecular mechanism of GLK, GNC and CGA1 in regulating hypocotyl elongation is not entirely clear. These need to be further investigated in combination with specific genetic material construction. It should be noted that additive relationships between HY5 and GLK, GNC, or CGA1 likely exist, in parallel with the fact of HY5 being upstream, given that HY5 acts to activate many transcription factor genes and functional genes the same time. In addition, GLK2 function has been standing out in multiple ways, partially recruited by and interact with HY5, and effectively compensates for HY5 function, so that it deserves targeted studies in order to guide the potential application of GLK2 and HY5 in crop enhancement.

### GLK2 functions downstream but partially independent of HY5 for a prominent role in seedling greening

HY5 and GLK both functions in regulating chlorophyll biosynthesis and chloroplast development, but targeted studies and clear conclusions are missing on whether or how the two are related. Studies have shown that PHYTOCHROME-INTERACTING FACTOR 4 (PIF4) in Arabidopsis binds to the promoters of *GLK1* and *GLK2* to inhibit their expression (Song et al., 2014), while HY5 antagonises PIFs to regulate the biosynthesis of photosynthetic pigments (Tolede-Ortiz et al., 2014). Besides, microarray studies showed that both phytochrome A and phytochrome B regulate GLK transcription (Tepperman et al., 2006). Considering that HY5 acts downstream of the photoreceptors, and is a high-level regulator (many HY5 targets are other transcription factors) of photomorphogenesis (Lee et al., 2007; Xu 2020), there may be a direct positive regulation of HY5 to *GLKs*. We provided experimental evidences in this work that HY5 indeed binds to the promoters of *GLK* genes and activates their expression (**Figure 3**). Similarly, the HY5 regulation of *GNC* and *CGA1* was also confirmed **(Supplemental Figure S3)**.

We have constructed double and triple mutants to study the functional relationships of these regulators. Decrease of chlorophyll content and related gene expression were observed when *hy5 glk2* mutant was compared to *glk2* mutant, or *hy5 glk1 glk2* mutant was compared to *glk1 glk2* mutant (**Figures 1 and 2**). Our transcriptome data indicate that, compared with HY5, GLK2 tends to positively affect more PSII and PSI related genes (**Figure 5D**), which was also supported by the stronger regulation of GLK2 on light harvesting genes than HY5 (**Figure 2B**). When overexpressing *GLK2* in the *hy5* mutant, the chlorophyll level increased significantly, although it was still lower than overexpressing *GLK2* in wild type (Liu et al., 2022). Our study clarifies the downstream relationship of GLKs to HY5, and implies that GLK2 contributes more intensively than HY5, to Arabidopsis seedling greening.

It has been reported that HY5 and GLK2 are involved in the greening of detached root in Arabidopsis, and that the GLK2 regulation of root greening depends to some extent on the presence of HY5 (Kobayashi et al., 2012a). We observed that the detached roots could still turn green in the absence of a single GLK or both GLKs, but not in the absence of HY5; similar trends were found in the detached roots of GNC, CGA1 and HY5 related mutants **(Supplemental Figure S7)**. The greening process is promoted by the accumulation of endogenous cytokinin in detached roots (Kobayashi et al., 2012a). In order to investigate the effect of HY5 and GLKs on cytokinin regulated leaf greening, we added 6-Benzylaminopurine (6-BA) during seedling greening of *hy5 glk* series mutants. Results showed that 6-BA treatment could promote the accumulation of chlorophyll during this process, with or without GLKs or HY5 **(Supplemental Figure S8)**. These results suggest that GLKs and HY5 play similar roles in response to cytokinin to promote chlorophyll biosynthesis in leaves, whereas HY5 plays a more significant role in detached roots.

GLKs seem to exert stronger regulation than GNC and CGA1 in seedling greening, as *glk* series mutants are much paler than *gnc* and *cga1* series mutants (**Figures 1 and 9; Supplemental Figures S1 and S2)**. No difference in chlorophyll content between *glk1 glk2 gnc cga1* quadruple mutant and *glk1 glk2* mutant was found, and *CGA1* overexpression in *glk1 glk2* mutant could not restore the chlorophyll content, while overexpression of *GLK1* partially restores the chlorophyll content in *gnc cga1* mutants (Bastakis et al., 2018). Zubo et al. suggested that GLK1, GLK2, GNC and CGA1 play both overlapping and independent roles in regulating chlorophyll biosynthesis and chloroplast development (Zubo et al., 2018). We further propose that GLK2 may play more prominent roles than GLK1, GNC and CGA1, downstream but partly independent of HY5, to regulate various targets during the de-etiolation process of photomorphogenesis.

### Differential participation of GLK and GNC in HY5 regulated hypocotyl elongation

The inhibition of hypocotyl elongation, opening of cotyledon, and chloroplast development are continuous and inseparable processes during seedling photomorphogenesis. Since HY5 inhibits hypocotyl elongation, we were also curious about the potential GLK and GNC regulation of hypocotyls. It has been mentioned that *glk1* seedlings displayed longer hypocotyls and less separated cotyledons (Martín et al., 2016; Alem et al., 2022). By systematically analysing the hypocotyl phenotype of mutants and overexpression lines, we certified that both GLK1 and GLK2 inhibit hypocotyl elongation, with additive effects on the function of HY5 (**Figure 4; Supplemental Figure S4)**. Transcriptomic sequencing and RT-qPCR verification showed that GLK2 itself could inhibit the expression of elongation genes, and different elongation genes exhibited different regulatory responses to GLK2 and HY5 (**Figures 4E and 5C**).

We noticed that the transcript levels of some elongation genes were reported lower in *gnc cga1* mutants than in the wild type (Xu et al., 2017), and the seedlings of the *quintuple* mutant of *GNC, GNL*, and *B-GATA* had shorter hypocotyl than wild type (Ranftl et al., 2016). In this work, we have confirmed that, contrary to GLK’s inhibitory effect, GNC and CGA1 promote hypocotyl elongation, and the mutation of GNC or CGA1 negatively affects the inhibition of hypocotyl lengths by HY5 (**Figures 9C and 9D; Supplemental Figure S5)**. Our results indicate that GNC could promote the expression of elongation genes to varying degrees, independently or on top of HY5 (**Figure 9E**). It should be pointed out that the same elongation gene maybe differentially regulated by HY5, GLKs, GNC or CGA1. In addition, the changes of elongation genes in *glk1 glk2, gnc cga1, hy5 glk1 glk2*, and *hy5 gnc cga1* mutants were also explored, showing similar additive effect as the *glk2* or *gnc* mutant on *hy5*.

### Significance of GLKs in the context of skotomorphogenesis to photomorphogenesis transition

Traits of photomorphogenesis are negatively regulated by the COP/DET/FUS complex in the dark, which mediates the ubiquitination and proteasome degradation of positive regulators such as HY5, HYH, LAF1, and HFR1 (Osterlund et al., 2000; Holm et al., 2002; Seo et al., 2003; Yang et al., 2005; Han et al., 2020; Qin et al., 2020). Interestingly, *det1* and *cop1* mutants exhibit some of the characteristics of photomorphogenesis during skotomorphogenesis, with shorter hypocotyls, open cotyledons, and expression of photosynthetic genes (Chory et al., 1989; Deng et al., 1992). These phenotypes should be related to the accumulation of positive regulators of photomorphogenesis in darkness, but the specific regulators and their influence on the phenotypes are not clear. It has been reported that long-term ABA treatment activates the activity of COP1, which then interacts with GLK1 protein to mediate its ubiquitination and degradation (Tokumaru et al., 2017; Lee et al., 2021). In this study we further showed that DET1 is likely responsible for the protein degradation of both GLK1 and GLK2 in Arabidopsis seedlings (**Figure 8A; Supplemental Figure S9)**.

Fonseca’s and Rubio’s groups showed that HY5 protein was more stable in the *det1* mutant, and demonstrated that DET1 associated with COP1 *in vivo*, enhanced COP1-HY5 interaction, thus influenced COP1 function in the regulation of HY5 protein (Cañibano et al., 2021). Since HY5 regulates *GLK* expression, it is possible that GLK protein accumulated more in the dark-grown *det1* mutant because HY5 protein is more stable. For this reason, we compared the *GLK1* and *GLK2* transcript levels in etiolated wild type, *det1, hy5* and *det1 hy5* double mutant. The result showed increased *GLK1* and *GLK2* transcript levels in etiolated seedlings of *det1* mutant than in that of wild type (**Figure 8B**), which is likely related with enhanced HY5 protein stability in *det1* mutant. Interestingly, *det1 hy5* double mutant showed *GLK1* transcript level similar to that in *hy5* mutant (lower than in wild type), while it showed *GLK2* transcript level similar to that in *det1* mutant (**Figure 8B**). Considering the similar expression of *GLK2* between WT and *hy5* in the dark and at 6h of de-etiolation but compromised *GLK2* expression in *hy5* after 12 h of greening (**Figure 1D**), it is likely that the regulation of *GLK2* expression by HY5 is more light dependent, while the regulation of *GLK1* expression by HY5 is already effective in etiolated seedlings. Therefore, the increased GLK1 protein content in dark-grown *det1* mutant was also HY5 dependent, while the increased GLK2 protein may be mainly resulted from impaired degradation. It is also possible that GLKs could induce their own expression.

As for GLK2, we have constructed *det1 glk2* mutant in Arabidopsis to compare with *det1* mutant during skotomorphogenesis. The overall number of extended thylakoid lamellae and expression of photosystem genes were found markedly reduced in *det1 glk2* compared with *det1* mutant (**Figures 7B and 8C**). In fact, both GLK2 and GLK1 contribute significantly to the expression of photosystem genes, not only in the light but also in dark-grown seedlings, while GNC and CGA1 seem to contribute less than GLKs (**Figures 8D and 8E**). Thus, the appropriate function of GLKs, GNC and CGA1 may be regulated at protein level by DET1 in etiolated seedlings, while upon light HY5 takes the transcriptional control and at least partly coordinates their functions, in a regulation cascade of light, HY5, and GLKs, GNC or CGA1.

We constructed a working model as shown in **Figure 10**. In dark-grown Arabidopsis seedlings, GLKs, GNC, CGA1 and HY5 proteins are subjected to the DET1 mediated degradation. Photosynthetic genes are inactive and plastid inner membranes stay at prolamellar body status. After exposure to light, DET1 activity is inhibited and GLK, GNC, CGA1 and HY5 proteins are able to accumulate. HY5 activates *GLK, GNC* and *CGA1* genes in the light, and they could act either independently, additively (on top of HY5), or cooperatively (HY5 likely interact with others) to regulate downstream gene expression and promote photomorphogenesis. GLKs are master regulators of chloroplast development and able to inhibit hypocotyl elongation independent of HY5. GNC and CGA1 are less dominant in chloroplast development and able to promote hypocotyl elongation. These could be the compensation for the relatively weak regulation of HY5 on chloroplast development, and a sensible way to orchestrate light induced seedling development, by recruiting multitasking yet complementary regulators. Our study systematically reveals the new function of GLKs, GNC and CGA1 regulating hypocotyl length under light, refines the local network of HY5, GLKs, GNC and CGA1, and provides new perspective for better understanding the transition between dark and light-grown Arabidopsis seedlings.

**Figure 10.**
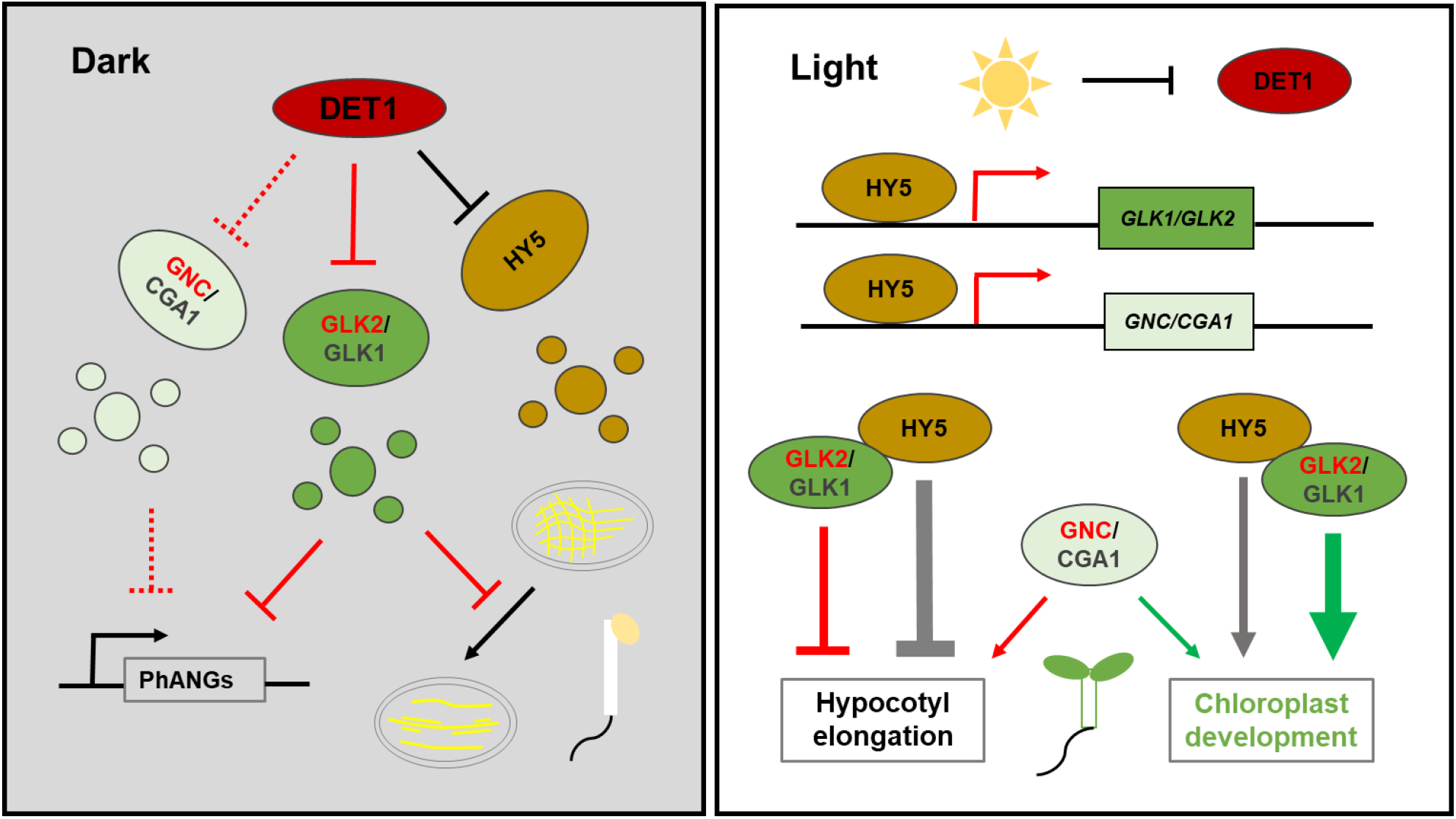
A working model depicting GLKs, GNC/CGA1, and HY5 in the regulation of skotomorphogenesis and photomorphogenesis. GLKs, GNC, CGA1 and HY5 undergo DET1-mediated degradation in the dark, so that the expression of photosynthesis associated nuclear genes (PhANGs), and the transformation of prolamellar body towards extended lamellae are inhibited. Light inhibits DET1 activity, and HY5 binds the promoter of *GLKs, GNC* and *CGA1*, inducing their activities to promote chloroplast development. The protein interactions between HY5 and GLKs are shown. GLKs inhibit hypocotyl elongation while GNC and CGA1 promote hypocotyl elongation. Thick green arrow indicates strong promotion of chloroplast development by GLKs. Thick gray line indicates strong inhibition of hypocotyl elongation by HY5. Thinner arrows indicate less strong regulation. Dashed lines in the dark indicate GNC/CGA1 related regulations that pending to be further validated.

## Materials and methods

### Plant materials and growth conditions

Arabidopsis mutants used in this study were in the Columbia (Col-0) background. The *glk1, glk2, glk1 glk2, gnc, cga1, gnc cga1, hy5*, and *det1* mutants (Chory et al., 1989; Oyama et al., 1997; Fitter et al., 2002; Chiang et al., 2012), as well as *35S:GLK1 glk1 glk2, 35S:GLK2 glk1 glk2, 35S:GNC*, and *35S:CGA1* overexpression plants (Waters et al., 2008; Chiang et al., 2012) were described previously. Other mutants described in this study were created by crossing from the above materials. Surface-sterilized seeds were sown onto half-strength Murashige and Skoog (MS) medium, which contained 1% sucrose and 0.8% agar, and were cold-treated at 4°C in darkness for 3 days to ensure synchronized germination, followed by growing in dark or light (100 μmol/m^2^/s) chambers maintained at 22°C.

### Chlorophyll measurements

Total chlorophylls were extracted by homogenizing the seedlings in 80% acetone, and leaving the solutions at 4°C for overnight incubation. The absorbance was detected at wavelength OD_652_ with a spectrophotometer. Chlorophyll concentration Ca+b was calculated by OD_652_/34.5 (mg/mL), and then converted according to the fresh weight of the material (Arnon, 1949; Zhao et al., 2016).

### Hypocotyl length measurement

In order to measure the hypocotyl length of seedlings, seeds were sown on plates and stratified at 4°C in darkness for 3 d, and then kept in white light for 4 h in order to induce uniform germination. The seeds were then transferred to white, blue, red light, or dark conditions and incubated at 22°C for 4 d. Using ImageJ software to measure the hypocotyl length after photographing the seedlings.

### Transmission electron microscopy

Leaf sections of 2 mm size were cut from the seedlings and fixed in 2.5% glutaraldehyde, with low vacuum applied for 60 min to assist fixation. Post fixation in osmium tetroxide, embedding in Spurr’s resin, and other steps were performed by the TEM platform in Center for Excellence in Molecular Plant Sciences, following standard procedures. The ultra-thin sections were imaged at 80 KV with a Hitachi H-7650 transmission electron microscope.

### Quantitative RT-PCR

Total RNA was extracted from Col-0 and mutant seedlings using an RNAiso Plus kit (Takara). Complementary DNAs (cDNA) were synthesized from 2 μg of total RNA using a Prime Script RT Reagent Kit, with a genomic DNA Eraser (Yeasen). Then, cDNA was subjected to RT-qPCR assays, using the Step One Plus RT-PCR detection system (Applied Biosystems) and SYBR Green PCR Master Mix (Yeasen). PCR was performed in triplicate for each sample, and the transcript levels were normalized to that of *ACT7* gene.

### Immunoblot analysis

Seedlings were homogenized in a protein extraction buffer containing 100 mM Tris (pH 6.8), 10% Glycerol, 0.5% SDS, 0.1% Triton, 5 mM EDTA, 0.01 M DTT, and 1 × complete protease inhibitor cocktail (Roche). Equal amount of proteins were loaded and separated on a 10% SDS-PAGE gel and transferred to polyvinylidene fluoride membrane. The membrane was blocked with 5% milk and incubated with primary antibody overnight at 4°C, washed four times with 1 × Phosphate Buffered Saline Tween-20 (PBST) for 5 minutes, and incubated with secondary antibody for 1 h at room temperature. After four washes with PBST for 5 minutes, signal was detected with ECL kit (Millipore).

### EMSA assay

Synthetic complementary oligonucleotides of *GLK1, GLK2, GNC*, and *CGA1* (or mutant probe with HY5 binding motif mutation) were obtained, and probes were PCR amplified using Cy5-labeled primers. For proteins, the coding sequence of HY5 was cloned into pCold-TF vector, expressed, and purified with Capturem™ His-Tagged Purification Maxiprep Kit (Takara). The binding reaction was performed in 20 μL binding buffer (10 mM Tris pH 8.0, 1 mM KCl, 4 mM MgCl_2_, 0.5mM DTT, 5% glycerol, 0.2 mM EDTA, and 0.01% BSA) using 15 nM probes and 200 ng proteins, and incubated at room temperature for 30 min. The reactions were resolved by electrophoresis in a 6% (v/v) native polyacrylamide gel at 4°C. Cy5-labeled DNA in the gel was scanned by an Amersham Typhoon 5 Biomolecular Imager.

### Dual-luciferase reporter system

Promoter sub-fragments of *GLK1, GLK2, GNC*, and *CGA1* were cloned into the pGreenII 0800-LUC vector to drive the firefly luciferase gene. p2GWY7-HY5-YFP was used as the effector construct. Arabidopsis mesophyll cell protoplasts were isolated and transfected as described previously (Yoo et al., 2007). We used a Dual Luciferase kit (Yeasen) for transient expression analysis to detect reporter activity. The Renilla luciferase gene, driven by the cauliflower mosaic virus 35S promoter, was used as an internal control. The ratio of LUC/REN was calculated as an indicator of the final transcriptional activity.

### Chromatin immunoprecipitation (ChIP)

The ChIP assay was operated as described by Xu et al. (2016). The 4-day-old Col-0 and *HA-HY5 hy5-215* transgenic seedlings grown under constant white light were used to isolate chromatin. 250-500 bp fragments were obtained by sonicating the resuspended chromatin at 4°C. The treated chromatin was immunoprecipitated, washed, and reverse cross linked. About 10% of sonicated but non-immunoprecipitated chromatin was reverse cross linked and used as an input DNA control. Monoclonal anti-HA antibody (ab9110, Abcam, 1:100 dilution) was used for immunoprecipitation. Both immunoprecipitated DNA and input DNA were analyzed by RT-PCR. The level of binding was calculated as the ratio between the IP and Input groups.

### *In vitro* pull-downs

The full-length coding sequences of *GLK1, GLK2, GNC*, and *CGA1* were cloned into pGEX-6P-1 vector, and the full-length coding sequences of *HY5* and *nDET1* (26-87 aa) were cloned into pCold-TF vector. Two mL of transformed bacteria cultures were grown over night, diluted 1:100 in the next morning, and grown for another 3 hours at 37°C. When the cells reached logarithmic phase, 0.1 mM IPTG was added to induce the expression of proteins at 37°C for 4 hours. Then 0.5 mL of each selected *E. coli* cultures was added in the same tube, re-suspended in lysis buffer (1 × PBS, pH 7.4, 1 × complete protease inhibitor cocktail), and the cells were broken with a sonicator for 3 times (30 s on and 60 s off) on ice. After the cell debris were removed by centrifugation at 18000 g for 30 min, 20 μL Glutathione Sepharose 4B was added to the mixture sample and rotated at 4°C for 6 hours. After 4 washes with PBST buffer (1 × PBS, pH 7.4, 0.1% Triton X-100), the pellet fraction was boiled in 5 × SDS protein loading buffer, and the input and pull-down prey proteins were detected by immunoblot using anti-His (M201, Takara, 1:3000 dilution) and anti-GST (G018, Abcam, 1:3000 dilution) antibodies respectively.

### Bi-luminescence complementation (BiLC) assay

GLK or HY5 was fused to the N- or C-terminus of firefly luciferase, and the constructs were transformed into agrobacterium (*Agrobacterium tumefaciens*) strain GV3101. Overnight cultures of agrobacteria were collected by centrifugation at 4000 g for 10 min, re-suspended in MES buffer (10 mM MES,10 mM MgCl_2_, and 100 mM acetosyringone), mixed with GV3101 colonies expressing pSoup-P19 to a final OD_600_ = 0.5, and incubated at room temperature for 3 h in the dark before infiltration. The agrobacterium suspension in a 1 mL syringe (without the metal needle) was carefully press-infiltrated onto healthy leaves of 3-week-old *N. benthamiana*. The infiltrated plants were returned to long-day conditions for 3 d. Leaves were infiltrated with luciferin solution and left for 10 min, before observing luciferase activity by imaging with a CCD camera (Tanon-5200, BioTanon, China).

### Co-immunoprecipitations (Co-IPs)

Arabidopsis mesophyll cell protoplasts isolated from two-week-old Col-0 plants grown in long-day conditions were used for Co-IP experiments. The protoplasts were transfected with a total of 10 μg DNA (p2GWY7-YFP, p2GWY7-HY5-YFP, and 35S-GLK2-2XFLAG-hcf) and incubated overnight. The transfected samples were spun for 2 min at 100 g, separated from the supernatant, and then homogenized in binding buffer (25 mM Tris-HCl, pH 7.5, 1% [v/v] Triton X-100, 150 mM NaCl, 1 mM EDTA, 10% [v/v] glycerol, and 1× protease inhibitor cocktail [Roche]) by rotating at 4 °C for 1 h. The insoluble material was removed by centrifugation at 13,000 g for 10 min at 4 °C, and the supernatant was mixed with 40 μL of ANTI-FLAG M2 affinity beads (Sigma-Aldrich). The mixtures were incubated at 4 °C for 6 h, and the beads were washed three times with washing buffer (25 mM Tris-HCl, pH 7.5, 0.5% [v/v] Triton X-100, 150 mM NaCl, 1 mM EDTA, and 10% [v/v] glycerol). We eluted the bound proteins from the affinity beads with 2× SDS-PAGE sample buffer and analyzed the eluates by immunoblotting with anti-GFP (SAB4301138, Sigma-Aldrich, USA, 1:1,000 dilution), anti-FLAG (F1804, Sigma-Aldrich, USA, 1:5,000 dilution), and anti-Actin (LF208, Epizyme Biotech, China, 1:3,000 dilution) antibodies to detect the target proteins.

### RNA-seq analysis

Total RNA was extracted from 4-d-old Col-0, *glk2, hy5*, and *hy5 glk2* seedlings grown in constant white light (100 µmol photons m^−2^ s^−1^). Sequencing libraries were generated with three independent biological replicates for each material, and the sequencing was performed using Illumina platform by Biomarker company (Beijing). Differentially expressed genes (DEGs) were identified using DESeq (Version1.18.0) (Anders et al., 2013).

### Accession numbers

Sequence information from this article can be found in The Arabidopsis Information Resource (TAIR) under the following accession numbers: *GLK1* (AT2G20570), *GLK2* (AT5G44190), *GNC* (AT5G56860), *CGA1* (AT4G26150), *DET1* (AT4G10180), *HY5* (At5G11260), *ACT7* (At5G09810), *HEMA1* (AT1G58290), *PORB* (AT4G27440), *PORC* (AT1G03630), *CAO* (AT1G44446), *LHCB1.2* (AT1G29910), *LHCB2.2* (AT2G05070), *LHCA1* (AT3G54890), *LHCA4* (AT3G47470), *EXPA5* (AT3G29030), *IAA19* (AT3G15540), *SAUR20* (AT5G18020), *PME16* (AT2G43050), *IAA7* (AT3G23050), and *XTH4* (AT2G06850). The references from which we selected the cell elongation genes are: Tatematsu et al., 2004; Schröder et al., 2009; Spartz et al., 2012; Zhou et al., 2013; Reed et al., 2018; Kushwah et al., 2020.

## Acknowledgments

We thank Prof. Jane Langdale for comments and suggestions on the work. We thank Xiao-Yan Gao and Zhi-Ping Zhang for technical support on the transmission electron microscopy. We are grateful to Prof. Hong-Hui Lin for kindly providing the AtGLK1 antibody; Chun-Guang Chen from ORIZYMES for generating the AtGLK2 antibody; Prof. G. Eric Schaller for gift of Arabidopsis GNC and CGA1 deficient mutants and overexpression lines.

