## Supplemental Figures for "HY5 regulates GLK transcription factors to orchestrate photomorphogenesis in *Arabidopsis thaliana*"

### Supplemental Figure S1

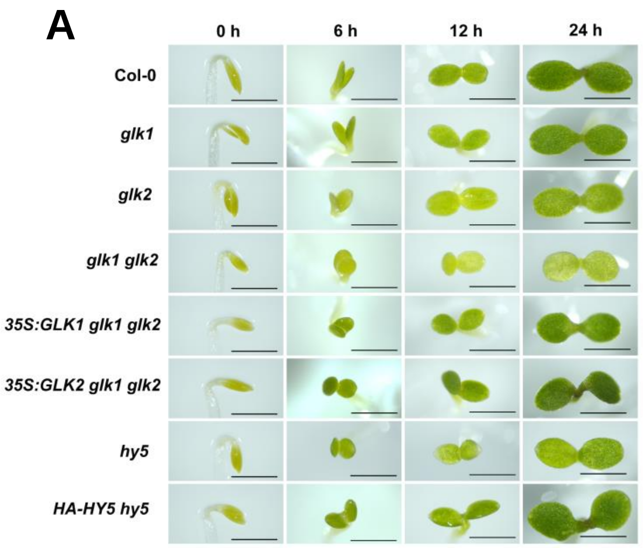


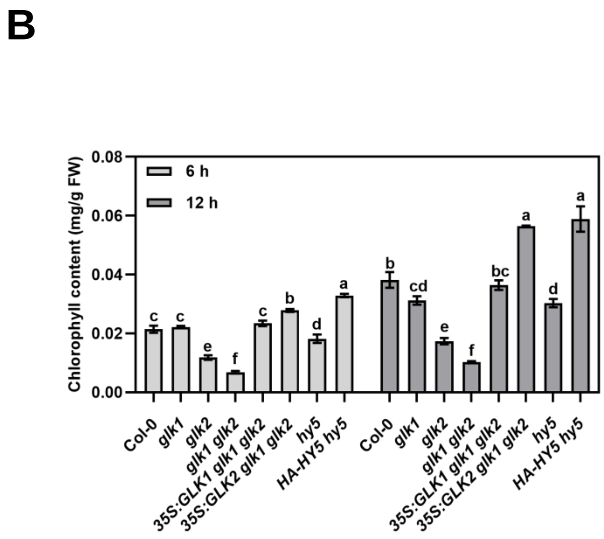


**Supplemental Figure S1.** HY5, GLK1 and GLK2 regulate the chlorophyll content of de-etiolating seedlings.
**(A)** Representative images of 4-day-old etiolated seedlings of Col-0, *glk1*, *glk2*, *glk1 glk2*, *35S:GLK1 glk1 glk2*, *35S:GLK2 glk1 glk2*, *hy5*, and *HA-HY5 hy5* during the transition from dark to light (100 μmol/m$^2$/s) conditions for 6 h, 12 h, and 24 h. Scale bars = 1 mm. **(B)** The whole seedling chlorophyll contents of 4-day-old etiolated seedlings from (A) during the transition from dark to light conditions for 6 h and 12 h. The data represent means ± SD (n=3) and letters above the bars indicate significant differences ($P$ < 0.05), as determined by one-way ANOVA with Turkey's HSD test.

### Supplemental Figure S2

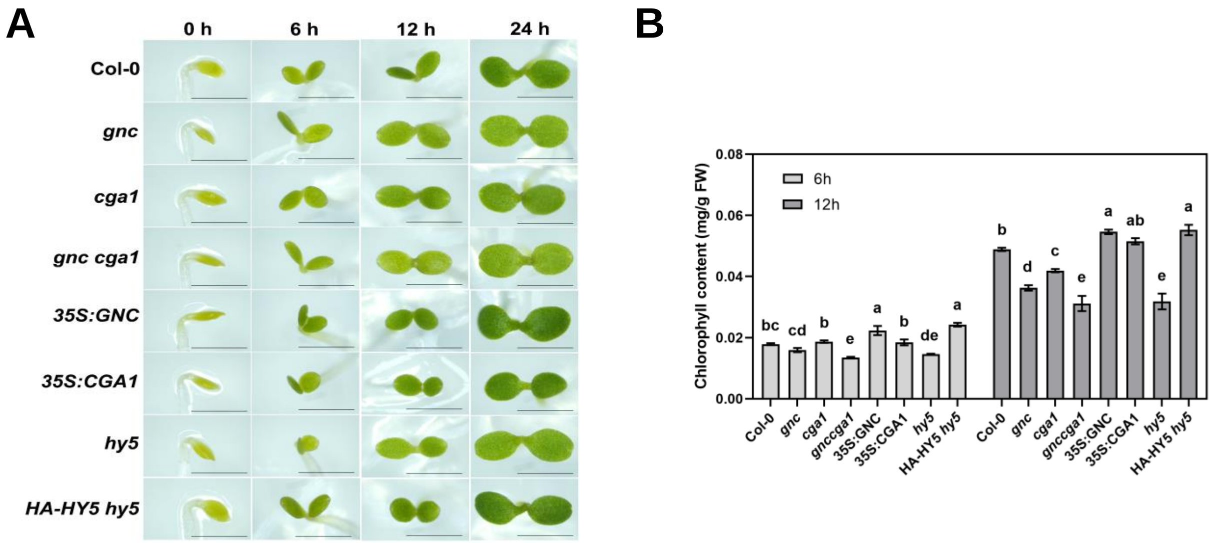


**Supplemental Figure S2.** HY5, GNC and CGA1 regulate the chlorophyll content of de-etiolating seedlings. **(A)** Representative images of 4-day-old etiolated seedlings of Col-0, *gnc*, *cga1*, *gnc cga1*, *35S:GNC*, *35S:CGA1*, *hy5*, and *HA-HY5 hy5* during the transition from dark to light (100 μmol/m$^2$/s) conditions for 6 h, 12 h, and 24 h. Scale bars = 1 mm. **(B)** The whole seedling chlorophyll contents of 4-day-old etiolated seedlings from (A) during the transition from dark to light conditions for 6 h and 12 h. The data represent means ± SD (n=3) and letters above the bars indicate significant differences ($P < 0.05$), as determined by one-way ANOVA with Turkey's HSD test.

### Supplemental Figure S3

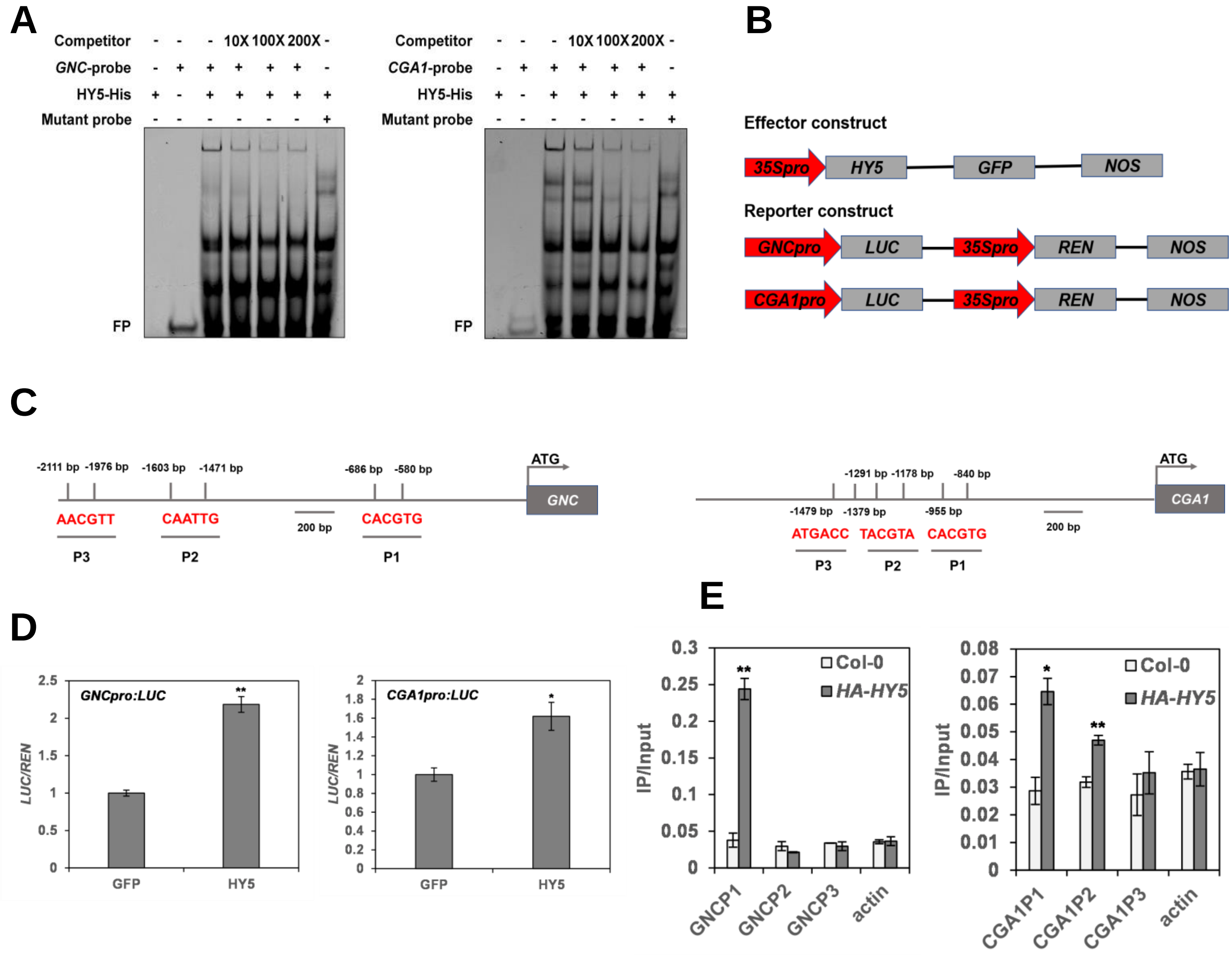


**Supplemental Figure S3.** Binding of HY5 to the promoters and transcriptional activation of *GNC* and *CGA1*.
**(A)** EMSA showing HY5 binds to the sub-fragments of promoters of *GNC* and *CGA1 in vitro*. "-" and "+" indicate the absence and presence of corresponding probes or proteins. FP means free probe. **(B)** Schematic structures of effector and reporter constructs used in dual-luciferase (LUC) reporter system. *REN*, renilla luciferase gene. **(C)** Illustration of *GNC* and *CGA1* promoter regions with the indicated positions of primers used in ChIP-qPCR. **(D)** Bar graphs showing HY5 induces the activation of *GNCpro:LUC* and *CGA1pro:LUC*. The data represent means ± SD (n = 4), and asterisks indicate a significant difference compared with control (**P*<0.05, ***P*<0.01, paired samples *t*-test). **(E)** Bar graphs showing ChIP-qPCR assays that HY5 associates with the promoters *in vivo*. Col-0 material and *ACTIN* gene were used as negative controls. The data represent means ± SD (n = 3), and asterisks indicate a significant difference compared with Col-0 (**P*<0.05, ***P*<0.01, paired samples *t*-test).

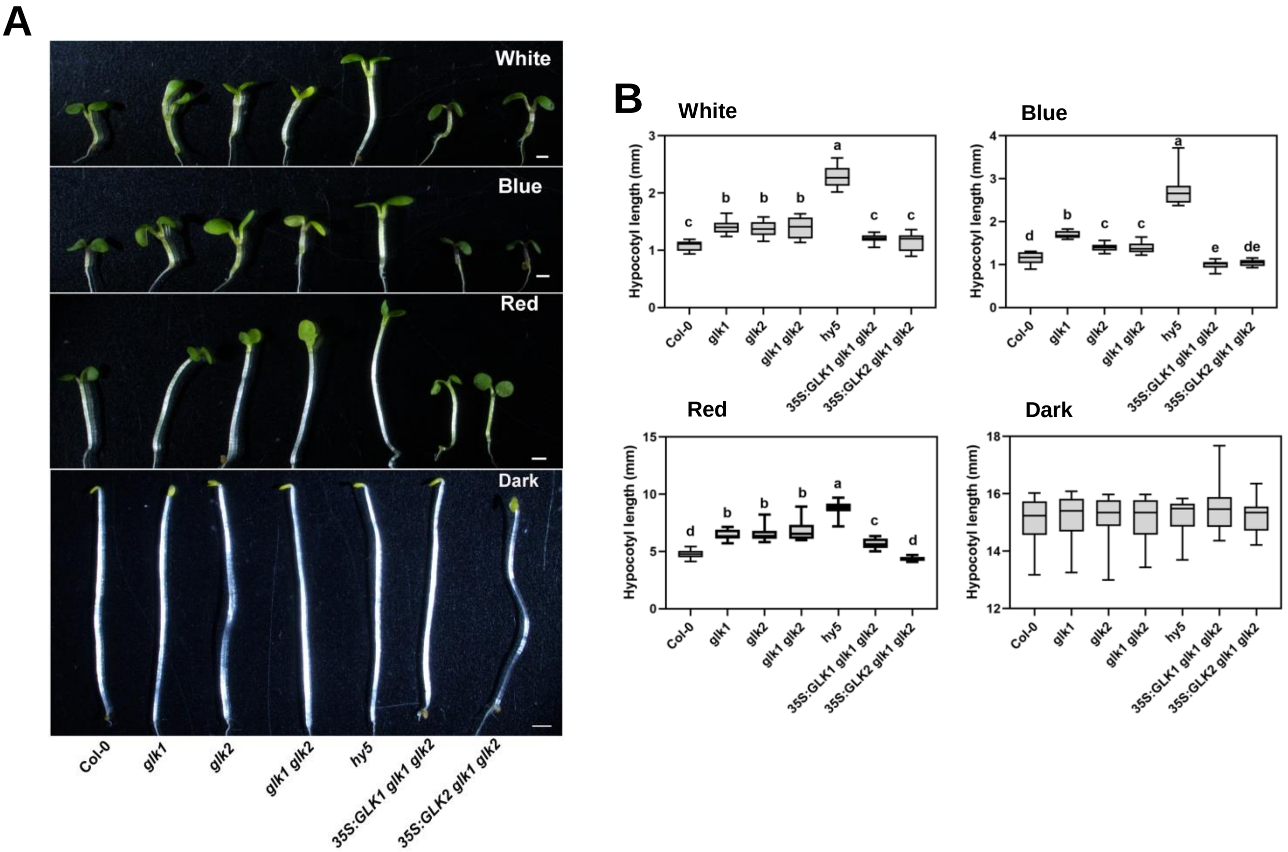


**Supplemental Figure S4.** GLK inhibits hypocotyl elongation. (Supports Figure 4)
**(A)** Hypocotyl phenotypes of 4-d-old Col-0, *glk1*, *glk2*, *glk1 glk2*, *hy5*, *35S:GLK1 glk1 glk2*, and *35S:GLK2 glk1 glk2* seedlings grown in continuous white (100 μmol/m$^2$/s), blue (60 μmol/m$^2$/s), red (90 μmol/m$^2$/s) light and dark conditions. Scale bar = 1 mm. **(B)** Quantification of hypocotyl lengths indicated in (A). The data represent means ± SD (n ≥18) and letters above the bars indicate significant differences ($P$ < 0.05), as determined by one-way ANOVA with Turkey's HSD test. The experiments were performed three times with similar results.

### Supplemental Figure S5

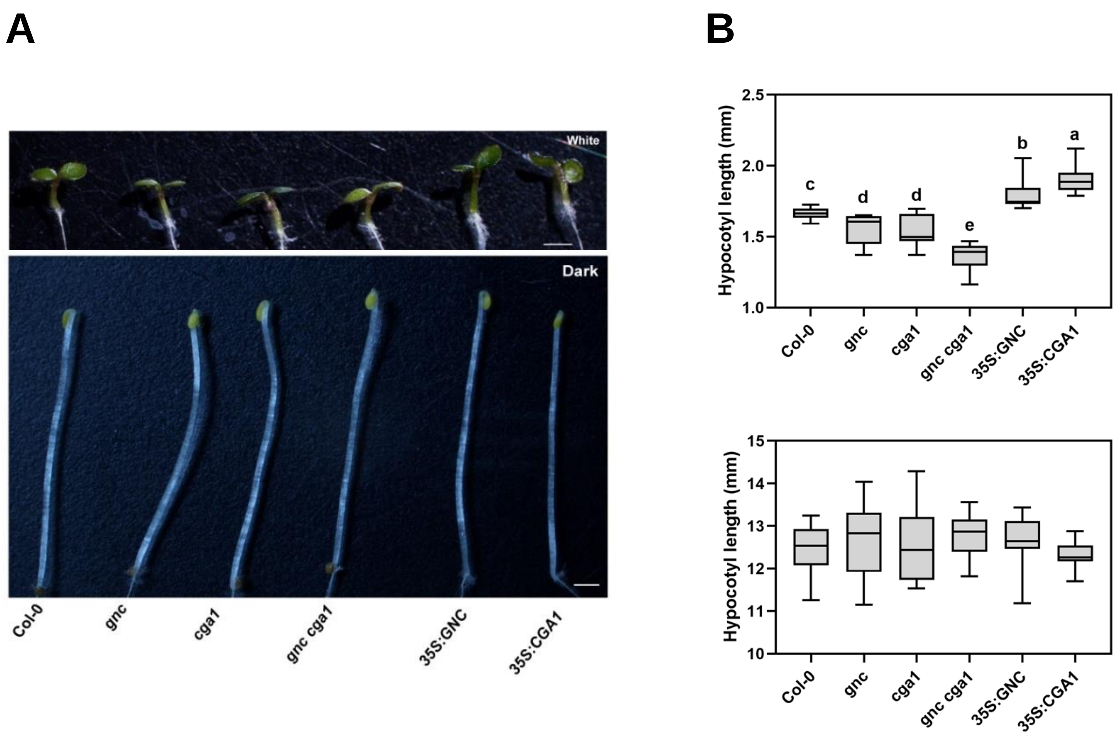


**Supplemental Figure S5.** GNC and CGA1 promote hypocotyl elongation.

**(A)** Hypocotyl phenotypes of 4-d-old Col-0, *gnc*, *cga1*, *gnc cga1*, *35S:GNC*, and *35S:CGA1* seedlings grown in continuous white light (100 μmol/m$^2$/s) and dark conditions. Scale bar = 1 mm. **(B)** Quantification of hypocotyl lengths indicated in (A). The data represent means ± SD (n ≥15) and letters above the bars indicate significant differences ($P$ < 0.05), as determined by one-way ANOVA with Turkey’s HSD test. The experiments were performed three times with similar results.

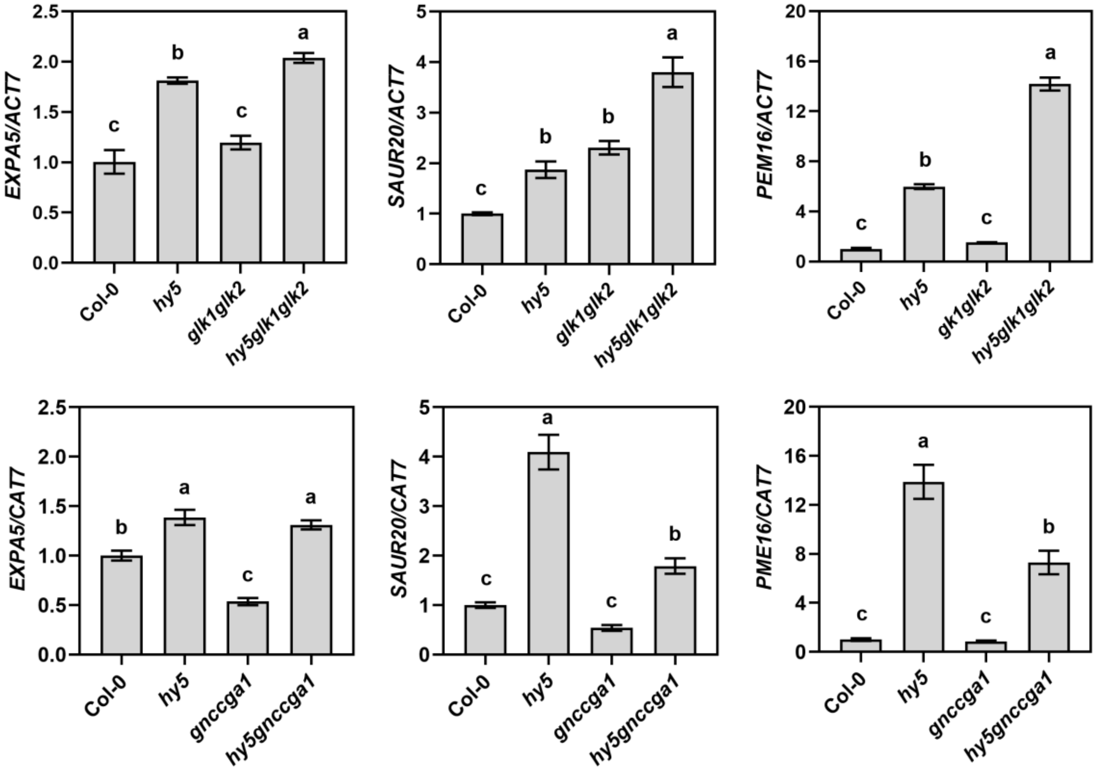


**Supplemental Figure S6.** Expression validation of elongation genes.
RT-qPCR analysis of cell elongation genes expressed in 4-d-old Col-0, *hy5*, *glk1 glk2*, *gnc cga1*, *hy5 glk1 glk2*, and *hy5 gnc cga1* seedlings grown under continuous white light (100 µmol/m$^2$/s). The *ACT7* gene was used as internal control. The data represent 3 replicates and letters above the bars indicate significant differences ($P$ < 0.05), as determined by one-way ANOVA with Turkey's HSD test.

### Supplemental Figure S7

A

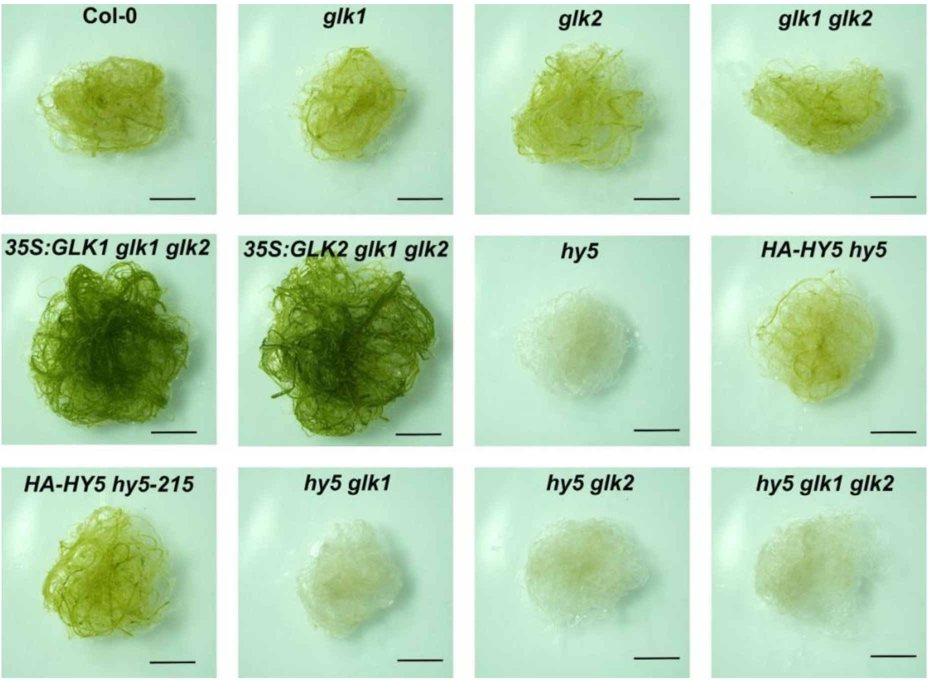


B

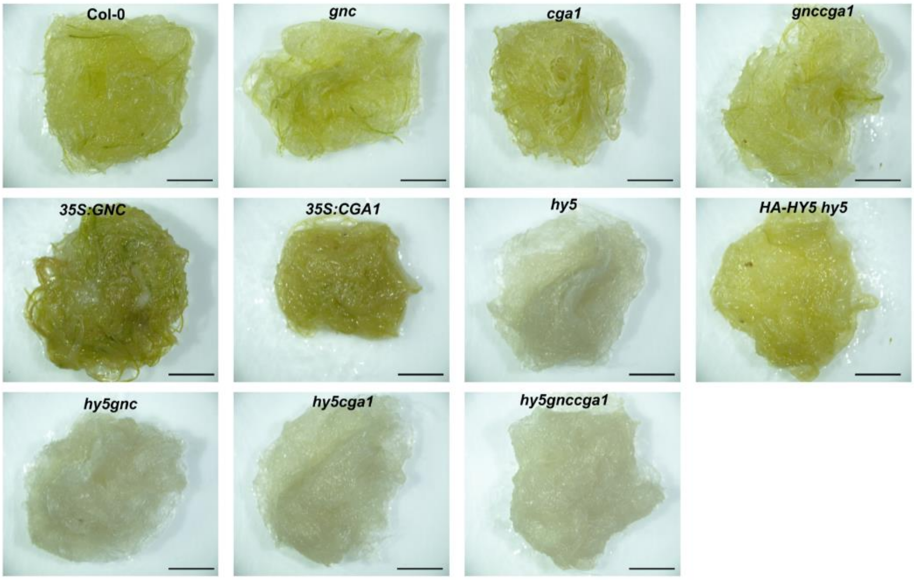


**Supplemental Figure S7.** HY5 but not GLKs (A) nor GNC and CGA1 (B) is crucial for the greening of detached roots.

Root samples were detached from 7-d-old seedlings, and then cultured on hormone-free MS medium for 14 d in continuous white light (100 µmol/m$^2$/s) conditions. Scale bar = 5 mm.

### Supplemental Figure S8

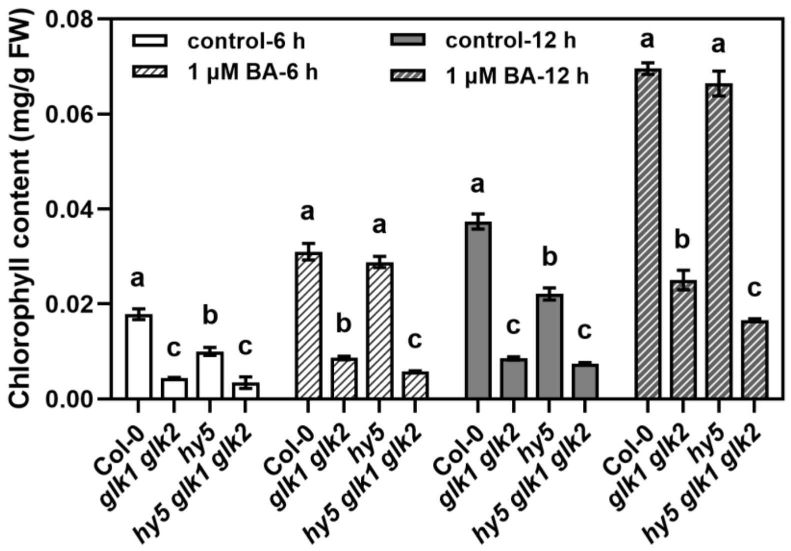


**Supplemental Figure S8.** 6-BA promotes the GLK and HY5 dependent chlorophyll content.
The whole seedling chlorophyll content of 4-day-old etiolated seedlings of Col-0, *glk1 glk2*, *hy5*, and *hy5 glk1 glk2* grown on medium with or without 6-BA during transition from dark to white light (100 μmol/m$^2$/s) conditions for 6 h and 12 h. The data represent means ± SD (n=3) and letters above the bars indicate significant differences ($P < 0.05$), as determined by one-way ANOVA with Turkey's HSD test. The experiments were performed three times with similar results.

### Supplemental Figure S9

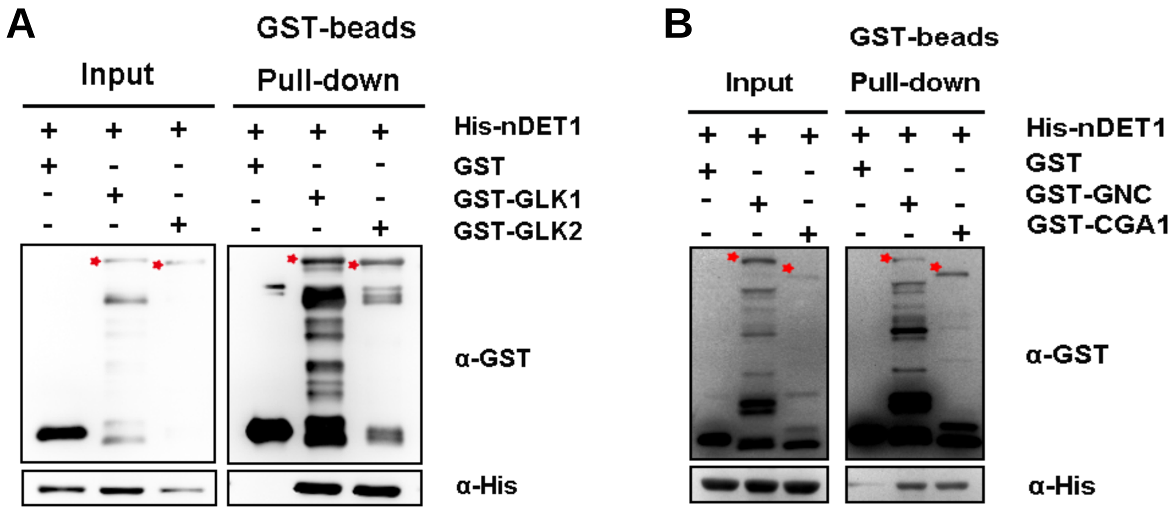


**Supplemental Figure S9.** *In vitro* evidence of DET1 interacting with GLK, GNC and CGA1 proteins. *In vitro* pulldown assays showing the interactions of GLK1, GLK2 (A), GNC and CGA1 (B) with DET1. GST-GLK1, GST-GLK2, GST-GNC, GST-CGA1 or GST proteins were used to pull down His-nDET1 protein using GST beads. Anti-GST and anti-His antibodies were used for immunoblot analysis. “-” and “+” indicate the absence and presence of corresponding proteins. The asterisks indicate GST-GLK1 (higher band), GST-GLK2 (lower band), GST-GNC (higher band) and GST-CGA1 (lower band) proteins.

### Supplemental Figure S10

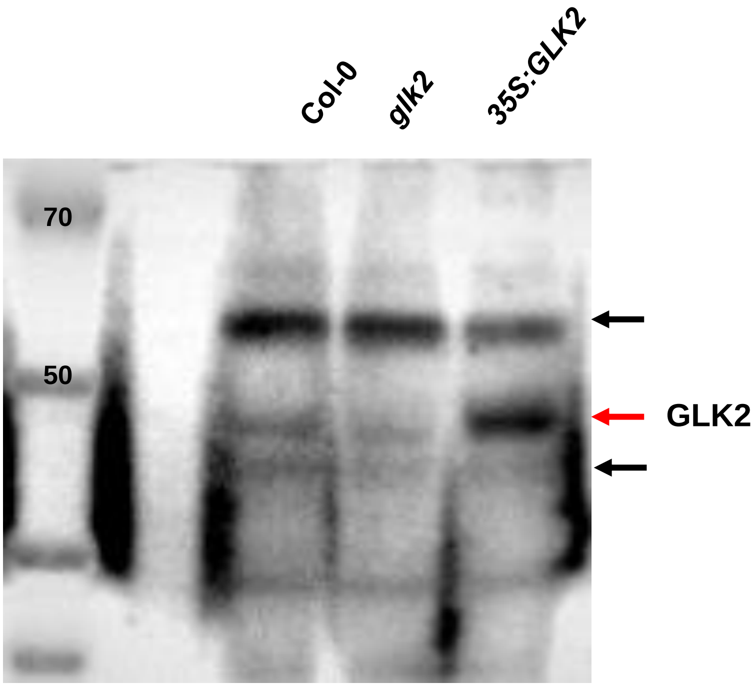


**Supplemental Figure S10.** Detection of GLK2 protein by anti-GLK2 antibody.
Total protein was extracted from 2-week-old Arabidopsis seedlings of Col-0, *glk2*, and *35S:GLK2* grown in growth chamber, resolved by SDS-PAGE, and probed with antibodies against GLK2. The red arrow indicates GLK2 protein and black arrows indicate nonspecific proteins.
